# Reconstructing developmental and disease progression with sample-level embeddings

**DOI:** 10.64898/2025.12.10.693462

**Authors:** Longda Jiang, Zhixin Cyrillus Tan, Isabella N. Grabski, Yuhan Hao, Nathan Nakatsuka, Sourav Sarkar, Anagha Shenoy, Rahul Satija

## Abstract

Single-cell genomics is transformative for characterizing cellular heterogeneity, but many translational questions require comparing samples rather than individual cells. Typical analyses compare “case” and “control” groups, ignoring sample-level variation within them. Here we present scSLIDE, a framework that transforms each sample’s single-cell data into a compact profile describing where its cells fall in high-dimensional space. By comparing these density profiles, scSLIDE calculates embeddings to characterization variation across samples. Applied to COVID-19 infection, Alzheimer’s disease, and zebrafish embryogenesis, we show that scSLIDE can be used to cluster patients, reconstruct sample-level disease trajectories, and identify coordinated cellular programs across samples. We discover independent axes of infection and severity, a molecular disease progression that aligns with pathology-based estimates of neurodegeneration, and embryonic “pseudostages” varying across and within timepoints. In each case, we demonstrate how case-control analyses collapse rich biological variation into binary labels, and how sample-level embedding represents a powerful analytical alternative.

## Introduction

Single-cell genomics has transformed our ability to characterize cellular identity, dynamics, and molecular circuitry at unprecedented resolution^1–3^. Most computational frameworks developed to date, have focused on individual cells as the fundamental unit of analysis, clustering them into discrete types and states^4,5^, constructing cell-level trajectories^6–8^, and transferring annotations from reference atlases^9–11^. While this cell-centric perspective has yielded critical biological insights, it falls short in many settings where the primary unit of interest is not a cell, but a sample.

In translational and clinical applications, researchers are often less concerned with describing the precise state of any single cell and more concerned with the distribution of cells within a sample. Case-control studies, patient stratification, and therapeutic response prediction all rely on detecting how cell types and states collectively differ across individuals^12–15^. For example, the clinically relevant questions focus not on which cluster individual cells belong to, but rather on how the cellular composition of a patient’s sample compares to others, whether an individual falls into a distinct disease cohort, and whether the cellular landscape is predictive of treatment outcome. Addressing such questions requires shifting from cell-level to sample-level embeddings to quantify similarities and discrepancies among samples and understand the drivers of sample-level heterogeneity.

In translational single-cell studies, the prevailing analytic paradigm has been to first assign cells to canonical types or states and then perform differential expression (DE) testing between health and disease groups within each cell type^16–18^. While widely adopted, this strategy suffers from two important limitations. First, it produces extensive lists of DE genes across many cell types, which are often difficult to interpret, prioritize, or integrate into coherent biological or clinical hypotheses^19^. Second, and more fundamentally, this framework implicitly assumes that all samples within a disease or control group are homogeneous representatives of their condition. This assumption is particularly problematic in the study of complex diseases, where heterogeneity is widespread and the disease labels themselves may be noisy or subjective^20–23^. Standard DE analyses average across these differences, potentially obscuring meaningful subgroups, disease trajectories, or predictors of therapeutic response.

Neurodegenerative diseases such as Alzheimer’s disease (AD) exemplify the challenges of analyzing complex conditions using traditional frameworks. Although AD has been the focus of extensive data generation^15,24,25^, even studies using similar profiling technologies may exhibit limited reproducibility when relying on case-control analyses^26–28^. This inconsistency can be driven by wide variation in pathological burden across patients that unfolds along continuous rather than discrete axes. Recent efforts have begun to move beyond binary labels, aiming to better characterize early-stage disease^29^ or to link specific cellular subpopulations to severity^30,31^. However, these analyses focus on cell states in isolation or are reliant on the abundance of predefined cell groups that are individually identified. New analytical frameworks that are capable of flexibly and reproducibly quantifying sample-level heterogeneity are therefore needed.

Here, we introduce a new computational framework, scSLIDE (single-cell Sample-Level Integration using Density Estimation), designed to directly characterize sample-level heterogeneity in single-cell data. scSLIDE leverages a semi-supervised dimensional reduction framework to embed cells into a latent space that robustly retains both their underlying type- and state-identity as well as phenotype-driven differences. Each sample is then represented as a density-based distribution of cellular states, yielding a sample-level representation that can be directly used for clustering, trajectory inference, and integrative analyses. We note that while powerful existing methods such as Milo^32^ also utilize density information, they do so to address a distinct problem: to perform binary “differential abundance” testing across conditions. In contrast, scSLIDE employs density to summarize the entire distribution of cellular states within each sample, uniquely enabling downstream analysis of sample-level heterogeneity.

We apply scSLIDE to three distinct settings, including viral infection^14,33^, neurodegenerative disease^25,30^, and zebrafish embryogenesis^34^ (Supplementary Table 1). In each case, scSLIDE uncovers sources of sample-level variation that are obscured by conventional, cell-centric approaches. We also find that the simple binarization of phenotypes into discrete “case” and “control” groups is inadequate, as samples often occupy diverse positions along continuous trajectories of cellular change. Leveraging this continuum substantially increases the statistical power, interpretability, and reproducibility of downstream analyses, such as the identification of disease-associated gene expression signatures. These results demonstrate that scSLIDE enables a shift from cell-centric to sample-level interpretation, offering a powerful and broadly applicable framework for single-cell studies.

## Results

### Overview of the scSLIDE method

To move beyond comparisons between individual cells and toward comparisons between samples, a computational framework must satisfy several requirements. First, it should capture cases where variation between samples arises from shifts in cell type abundance, such as expansion or depletion of specific cell subsets. Second, it should also detect differences within cell states, for example activation of inflammatory response programs in one or more cell types, and be capable of analyzing changes in cell type and cell state abundance simultaneously. Third, it must be sensitive enough to resolve subtle phenotypes, where relevant disease-associated signals constitute only a small fraction of the transcriptome. Finally, while the method can benefit from previously obtained sample metadata, it should not depend exclusively on it and must retain the ability to uncover previously unrecognized axes of variation.

The scSLIDE workflow is designed to address these requirements through three key steps (Figure 1a). As discussed in more detail below (with a full description in Supplementary Methods), we first embed cells into a latent space that preserves both broad cell type structure and finer-grained variation, combining unsupervised and metadata-aware representations through our previously introduced weighted nearest neighbor (WNN) approach^10^. Second, we estimate sample densities across this embedding by selecting a diverse set of “landmark cells”^35^ that span major types and states, and quantifying the density of each donor’s cells in the vicinity of each landmark. This procedure yields a compact sample-by-landmark matrix that summarizes sample-level heterogeneity and highlights disease-relevant differences. Third, we directly use this sample-level representation for downstream analyses, including clustering of samples, visualization of inter-sample relationships, trajectory inference, and identification of differential features associated with phenotypes of interest (Figure 1b). We describe each of these steps in detail below, and then apply and evaluate our framework across use-cases in immunology, neuroscience, and development.

**Figure 1.**
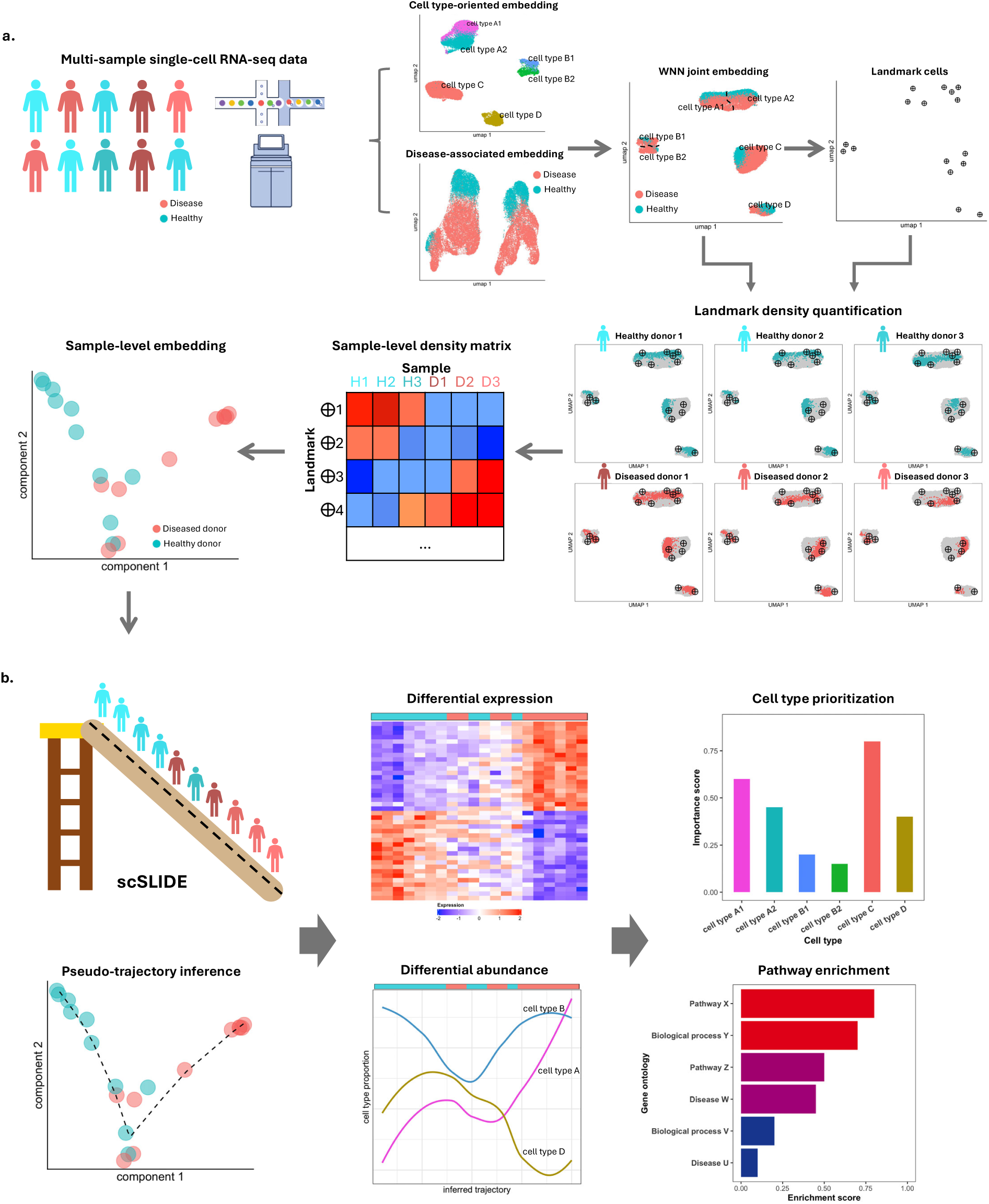
Overview of the scSLIDE framework for sample-level analysis of single-cell data. **(a)** scSLIDE converts multi-sample single-cell RNA-seq data into a sample-level representation. Cells are first embedded using a weighted nearest-neighbor (WNN) framework that integrates a cell-type-oriented (unsupervised) and a disease-associated (metadata-aware) embedding to preserve both cell-type resolution and phenotype-linked variation. A diverse set of landmark cells is then sketched to span major cell types and states, and sample-specific cell densities are quantified around each landmark to generate a sample-by-landmark density matrix. Finally, sample-level relationships (i.e. sample-level embedding) can be learned using dimensional reduction techniques. **(b)** The scSLIDE sample-level embedding enables a range of downstream analyses. Samples can be ordered along a pseudo-trajectory that represents continuous biological progression (illustrated as samples “sliding” down a trajectory of disease progression). This unified framework supports differential expression, differential abundance, cell-type prioritization, and pathway enrichment analyses, revealing molecular and compositional programs that drive sample-level phenotypes.

### Constructing a unified embedding of cell type and cell state variation

The first task in scSLIDE is to identify a low-dimensional space suitable for estimating cellular densities. This space must preserve high-resolution information on cell types while also capturing fine-scale differences in cellular state that may reflect key phenotypes or outcomes of interest, such as disease status, severity, or developmental timepoint. Generating this single-cell embedding via unsupervised analysis remains challenging, particularly as signals associated with disease-level variation can be weak and reflect only a small fraction of the transcriptome that is too subtle for unsupervised detection. Supervised dimensional reduction offers a powerful alternative, in particular, partial least squares (PLS) identifies low-dimensional projections that maximize covariance with outcome variables^36^. This approach is well-suited for separating cells based on known phenotypes but limits the possibility of discovering new sources of variation and comes at the cost of losing fine-grained information about cell types (Supplementary Figure 1a,b).

We reasoned that supervised and unsupervised strategies provide complementary perspectives, much like RNA and protein modalities in multimodal single-cell analysis. To integrate these perspectives, we adapt our previously introduced WNN framework, which unifies multiple embeddings into a single representation^10^. scSLIDE generates two intermediate representations. First, a “type”-focused embedding, which can be generated by integration tools^37–39^ or reference mapping methods^10,11,40^, prioritizes high-resolution cell type. Second, a “state”-focused embedding, generated with PLS, prioritizes separating cell states based on provided sample metadata (e.g., disease status). Combining these with WNN yields a single semi-supervised space that retains high-resolution cell type information, preserves the ability to discover novel sources of variation, and simultaneously prioritizes features associated with phenotypes of interest.

### Landmark-Based Quantification of Sample Densities

Once a suitable embedding has been constructed, the next step in scSLIDE is to estimate how each sample is distributed across this space. Inspired by methods developed to estimate sample density^35^, we identify a diverse set of landmark cells that serve as reference points and collectively span the cell types and states observed in the dataset. We utilize geometric sketching^41,42^ to ensure that landmarks cover both abundant and rare cellular states. In practice we select 5,000 landmarks per analysis but find that our results are robust across a range of this parameter (Supplementary Figure 1c). For each landmark, we estimate the density of cells from the sample that fall within its high-dimensional WNN local neighborhood. We accomplish this by first independently examining nearest-neighbor relationships between each cell and the full set of landmarks (i.e., identify the *k*-nearest landmarks for each cell) and subsequently aggregating the results across all cells within a sample (Supplementary Methods). We repeat this procedure for each sample individually, generating a “landmark abundance” matrix.

To account for differences in sample-level cell count and cell state frequency, we normalize the “landmark abundance” matrix using a chi-square style transformation. This approach highlights whether a sample is enriched or depleted for cells near a given landmark relative to what would be expected under independence. Specifically, we calculate the expected count for each entry based on the product of its row and column marginals and then scale the observed deviation by the square root of this expectation (Supplementary Methods). The resulting residuals capture relative density: positive values indicate that a sample has more cells than expected near a particular landmark, while negative values indicate a lower-than-expected density.

This normalized representation is a compact sample-level representation that fundamentally transforms the underlying single-cell measurements into a distributional profile for each sample, and emphasizes biologically meaningful shifts in composition or state. We call this output the “sample-level relative density matrix”, in which each column summarizes how a sample’s cells populate the landmark-defined state space. These vectors enable direct and quantitative comparisons between samples: cosine distances between columns reflect differences in composition and cellular state, much like expression-based measures are used to calculate distances between cells. Once in sample space, standard analyses, including clustering and trajectory inference, can be applied to identify disease subgroups, reconstruct sample-level progressions, and reveal coordinated biological changes across individuals. Thus, scSLIDE fundamentally restructures single-cell data into interpretable sample-level profiles that capture meaningful sample heterogeneity.

### scSLIDE identifies distinct axes of sample heterogeneity among COVID-19 patients

We first applied scSLIDE to a published single-cell dataset of COVID-19 infection^14^, which profiled immune responses in 78 COVID-19 infected individuals (spanning three severity categories) and 10 healthy controls. In total, the dataset comprises over 700,000 cells. We ran the full scSLIDE workflow, providing the PLS embedding with disease state and severity metadata (“mild”, “severe”, and “critical”) to help guide the representation. The resulting cell embedding retained high-resolution delineation of cell types and robustly separated cells from infected and control individuals (Figure 2a,b), from which we further generated the sample-level density matrix and performed normalization as described above.

**Figure 2.**
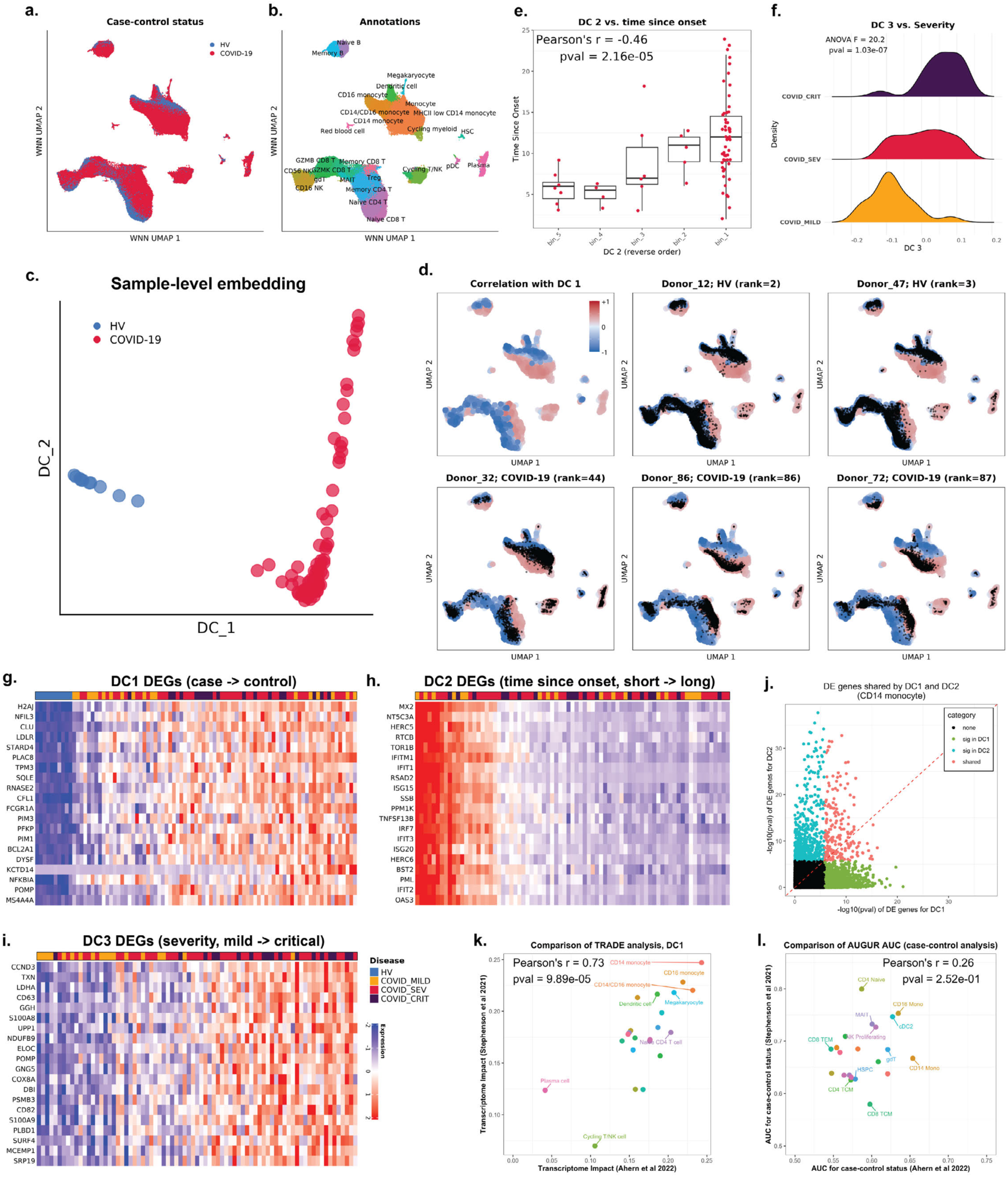
scSLIDE identifies distinct axes of sample heterogeneity among COVID-19 patients. **(a,b)** Uniform manifold approximation and projection (UMAP) of cell-level weighted nearest neighbor (WNN) embeddings from the COMBAT COVID-19 dataset after scSLIDE processing. **(a)** Cells colored by case–control status (HV, healthy volunteers; COVID-19, infected donors). **(b)** Cells colored by major immune cell annotations, showing preserved cell-type resolution. **(c)** Diffusion map embedding of sample-level relative density matrices identifies principal diffusion components (DCs), with DC1 separating cases from controls and DC2 capturing heterogeneity among infected donors. **(d)** Visualizing the density of cells from each individual sample. Each colored point represents a landmark, and is colored by Pearson correlation (red, positive; blue, negative) with DC1. Cells from five representative donors (black points) are overlaid onto the landmark map, illustrating how donor cells are distributed from samples across the DC1 gradient. Healthy donors primarily align with negatively correlated regions, whereas COVID-19 patients progressively expand into positively correlated areas, reflecting coordinated remodeling of immune cell states during infection. **(e)** DC2 correlates strongly with time since symptom onset (TSO) (Pearson r = –0.46, p = 2.16 × 10⁻⁵). **(f)** DC3 delineates a continuum of disease severity (ANOVA p = 1.03 × 10⁻⁷), ranging from mild to critical COVID-19 cases. **(g–i)** Heatmaps of representative differentially expressed (DE) genes whose donor-level pseudobulk expression correlates with DC1 (g), DC2 (h), and DC3 (i) in CD14 monocytes, ordered by diffusion component score. The legends for all three heatmaps are merged and shown in (i). **(j)** Limited overlap of DE genes between DC1 and DC2 indicates largely distinct transcriptional programs. **(k)** Comparison of transcriptome-wide impact (TI) scores from TRADE analysis across cell types shows strong reproducibility between independent datasets (Pearson r = 0.73). **(l)** Comparison of Augur-based case–control cell-type prioritization reveals lower reproducibility across datasets (Pearson r = 0.26).

To characterize sample heterogeneity, we applied diffusion maps^7^, a dimensional reduction method well-suited for capturing continuous trajectories, to the sample-level relative density matrix. The first diffusion component (DC1) perfectly separated cases from control samples (Figure 2c). Visualizing cells from each sample on the WNN UMAP, and arranging samples according to DC1, highlighted coordinated shifts in cellular state across multiple immune cell types (Figure 2d; Supplementary Figure 2a), consistent with broad inflammatory and antibody-generation responses induced by infection.

The second diffusion component (DC2) exhibited minimal variation among controls but captured substantial heterogeneity within the infected cohort (Figure 2c). Exploration of sample metadata revealed that DC2 position correlated strongly with time since disease onset (TSO) (Figure 2e, Supplementary Figure 2b), though not with disease severity (Supplementary Figure 2c), uncovering a clear source of temporal heterogeneity that varied across samples both phenotypically and molecularly. Notably, we did not provide TSO information to the scSLIDE workflow, but it still identified this axis of variation. Finally, the third diffusion component (DC3) stratified patients according to disease severity (Figure 2f). Unlike DC1, this component did not reveal discrete groups but instead arranged infected samples along a continuum from mild to critical outcomes (Figure 2f, Supplementary Figure 2d). We also observed a lack of correlation between sample position on the severity and TSO-driven axes (Supplementary Figure 2e), demonstrating how scSLIDE can disentangle independent axes of sample variation that would be blended together in traditional case-control analyses.

We next sought to determine which genes were associated with each DC. To do this, we focused on individual cell types and ordered donors according to their positions along a given diffusion component. For each gene, we then tested whether its expression correlated with the DC trajectory using a negative binomial generalized linear model (NB-GLM), with donor-level pseudobulk counts as the response and diffusion component score as a continuous predictor (Supplementary Methods). By repeating this trajectory-based DE test for each cell type, we could identify genes whose expression systematically increased or decreased along each axis of variation, rather than relying solely on discrete case-control labels.

When exploring gene expression changes, we focused first on CD14 monocytes, a myeloid cell type known to exhibit a strong transcriptional response to viral infection ^43^. For DC1, which cleanly divides samples into case and control groups, the NB-GLM test returned gene sets characterizing broad inflammatory and antiviral programs, representing a general signature of infection (Figure 2g, Supplementary Figure 2f). Since an axis representing a binary subdivision is well-represented by a case/control split, these genes closely mirrored those that were identified by conventional DE testing (Supplementary Figure 3a).

The continuous nature of DC2 and DC3 allowed NB-GLM to detect broad quantitative gene associations, revealing thousands of genes with significant associations that varied quantitatively along axes delineating additional variation within infected samples (Supplementary Table 2). DC2 was dominated by interferon-β–stimulated genes, reflecting interferon responses that were rapidly induced at disease onset (independent of severity) but that subsequently dampened over time (Figure 2h, Supplementary Figure 2g). Top DC3 genes were unrelated to interferon response but instead highlighted enrichment of genes involved in neutrophil degranulation, a pathway that has been specifically linked to myeloid responses in severe COVID-19^44,45^ (Figure 2i, Supplementary Figure 2h). We emphasize that the programs associated with each component were generally non-overlapping (Figure 2j, Supplementary Figure 3b) but would be indistinguishable in a traditional case-control analysis.

To identify which cell types are most transcriptionally affected for each of these components, we incorporated a method to quantify the *transcriptome-wide impact* (TI) of each DC on each cell type. This approach leverages the gene-level coefficients from our NB-GLM analysis, combined with the recently introduced TRADE model^46^ for assessing TI, to evaluate how strongly each axis perturbs the overall transcriptional landscape of a given population (Supplementary Methods). All major immune cell types showed measurable impact across the three components, but myeloid cells (in particular CD14 monocytes, CD16 monocytes and dendritic cells) were most strongly affected (Supplementary Figure 3c-e). Amongst lymphoid populations, Naive CD4 T cells showed elevated TI along DC1 (Supplementary Figure 3c), while Mucosal Invariant T (MAIT) and memory CD4 T cells exhibited stronger transcriptomic shifts along DC2 and DC3 respectively (Supplementary Figure 3d,e).

To assess the robustness of these findings, we applied our methods to an independent COVID-19 dataset^33^ (12 healthy donors and 43 patients, see Supplementary Methods) (Supplementary Figure 4a-d). When running scSLIDE on this new data, the first two diffusion components reflected case-control status and TSO (Supplementary Figure 4b,e), independently reproducing our findings from the Ahern et al dataset^14^. DC3 delineated mild from more serious COVID cases, but did not robustly separate moderate, critical, and severe patients, likely due to the reduced number of patients in this dataset (Supplementary Figure 4f). For DC1 and DC2, top DE genes associated with both components were also reproducible across independent datasets (Supplementary Figure 4g-j and Supplementary Figure 5a,b), as were our TI-based estimates of the most transcriptomically affected cell types (Figure 2k, Supplementary Figure 3f).

As cell-type prioritization is a key step for interpreting clinical data, we compared the results of our TI-based workflow with Augur^47^, a widely used tool for quantitatively identifying and ranking disease-relevant cell types based on single-cell case-control status. We found that Augur exhibited lower reproducibility across datasets, for example, prioritizing Naive CD4 T cells above all myeloid cell types in one dataset but not the other (Figure 2l). While powerful, Augur’s approach is limited to binary comparisons of a single-variable, and unlike scSLIDE, can therefore be confounded by additional sources of sample heterogeneity within either group. Together, these analyses demonstrate that scSLIDE provides interpretable and reproducible identification of gene programs, and prioritization of cell types linked to sample-level axes of variation.

### scSLIDE recovers a pathology-validated severity trajectory from Alzheimer’s snRNA-seq data

We next applied our framework to a more challenging setting: Alzheimer’s disease (AD), a condition marked by substantial heterogeneity in both clinical presentation and pathological features^48^. We first applied scSLIDE to a published SEA-AD single-nucleus RNA-seq dataset (10x 3’ snRNA-seq) generated by the Allen Institute^30^, which profiled cortical brain tissue across 75 individuals with varying degrees of Alzheimer’s disease (AD) pathology, along with 14 control samples. We also independently examined a second published snRNA-seq dataset from the Psych-AD consortium (150 cases, 149 controls, 10x 3’ snRNA-seq profiling)^25^.

We first applied a standard case-control DE analysis pipeline, where cells are “pseudo-bulked” by donor and cell type, followed by DE comparisons of binary cases vs. control groups across matched cell types (Supplementary Methods). To assess the robustness and reproducibility of this approach, we first analyzed Psych-AD (299 individuals) as a discovery cohort, and then separately analyzed SEA-AD (89 individuals) as a replication cohort. We then tested whether the significant DE genes identified in Psych-AD could be nominally replicated in SEA-AD (Supplementary Methods). Consistent with previous studies^27^, we found that only a low number (median: 17) and percentage (median: 17.8%) of the genes successfully replicated (Figure 3a, Supplementary Figure 6). The implicit assumption in DE workflows that case and control samples represent homogeneous groups could in principle help explain the lack of reproducibility across studies. We therefore examined if scSLIDE, which can more flexibly discover and account for continuous sources of heterogeneity, could improve the power and reproducibility for discovery of AD-associated gene modules.

**Figure 3.**
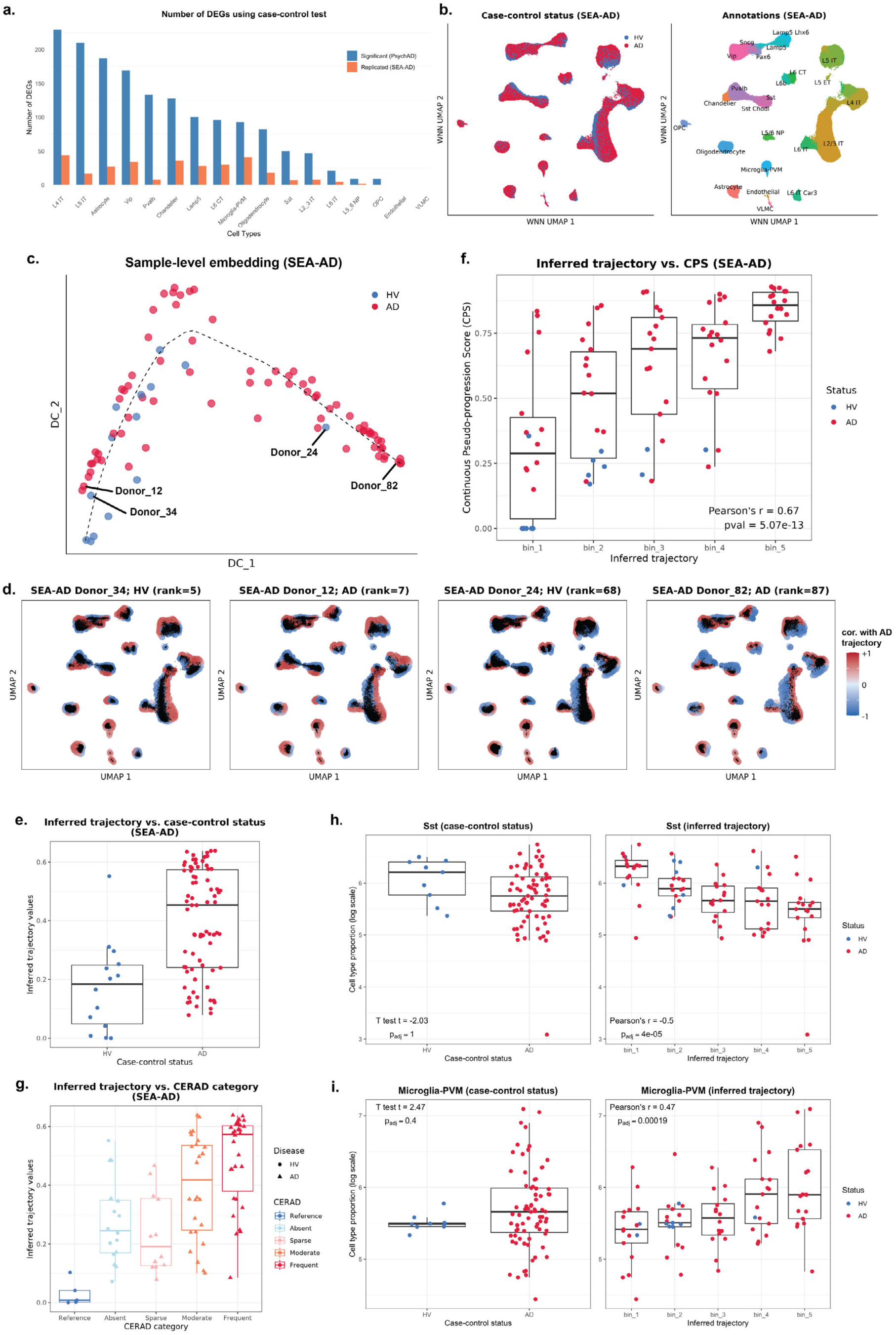
scSLIDE reconstructs a pathology-validated trajectory of Alzheimer’s disease progression. **(a)** Numbers of differentially expressed genes (DEGs) per cell type detected by binary case–control analysis in the Psych-AD discovery cohort (blue), with nominally replicated genes in the SEA-AD cohort (orange). **(b)** Cell-level weighted nearest-neighbor (WNN) embedding of the SEA-AD dataset colored by case–control status (left) (HV, healthy volunteers; AD, Alzheimer’s disease) and by annotated cell types (right). **(c)** Diffusion-map embedding of the sample-level relative density matrix reveals a continuous trajectory of AD progression rather than discrete case–control separation. A principal curve fitted through the first two diffusion components defines an inferred AD pseudo-trajectory across donors. **(d)** Landmark WNN UMAP showing landmark-trajectory correlation (red, positive; blue, negative), with cells from four representative donors overlaid, illustrating AD variability along the inferred pseudo-trajectory. **(e)** Donor positions along the trajectory relative to case–control status show extensive heterogeneity within clinically diagnosed AD samples. **(f)** The inferred trajectory significantly correlates with independently derived continuous pseudoprogression scores (CPS) reflecting neuropathological burden (Pearson r=0.67, p-value=5.07×10^−13^). **(g)** Donor positions along the trajectory stratified by CERAD plaque burden confirm association with pathological severity. **(h,i)** Cell-type abundance shifts along the inferred trajectory compared with binary case–control analysis. *SSTe* interneurons (h) decrease and microglia (i) increase progressively along the trajectory, consistent with neuronal loss and reactive gliosis in AD.

We first applied scSLIDE to the SEA-AD dataset, generating an intermediate cell-level embedding (Figure 3b) using WNN, and a sample-level embedding (Figure 3c) by running diffusion maps on the sample-level relative density matrix. We found that diffusion map components 1 and 2 exhibited continuous rather than discrete case-control separation (Figure 3c). We therefore fitted a joint principal curve^49^ to these components to infer a pseudo-trajectory of disease progression across donors (Figure 3c) (Supplementary Methods). This trajectory-based approach revealed significant intra-group heterogeneity that would be missed by traditional case-control comparisons. Importantly, variation along the cellular trajectory reflected underlying biological heterogeneity that was not captured by clinical diagnosis alone. In particular, while some diagnosed AD patients exhibit clear and coordinated shifts in cellular density across multiple cell types (i.e. Donor_82), at the other end of the trajectory AD patients (i.e. Donor_12) exhibited cellular density more similar to controls (Figure 3c,d, Supplementary Figure 7a).

While suggestive of extensive AD-heterogeneity among the patients (Figure 3e), scSLIDE’s trajectory represents a computational prediction that requires independent validation. In the SEA-AD study, each sample received a detailed neuropathological assessment followed by Bayesian modeling to assign each sample a continuous pseudoprogression score (CPS) that reflects disease severity. The CPS score reflects known features of disease burden including the abundance of pTau-bearing neurons, amyloid-Beta plaques, and signs of astrogliosis^30^. Since it does not utilize snRNA-seq data, and was not provided to the scSLIDE model, it provides an opportunity for independent validation of scSLIDE’s identified trajectory. Confirming this prediction, we identified a clear association between the scSLIDE trajectory and CPS score (Pearson’s r=0.67, p-value=5.07×10^−13^) (Figure 3f), confirming the biological relevance of this axis of heterogeneity. This validation revealed the full spectrum of AD heterogeneity: clinically diagnosed AD cases in the earliest progression decile had lower CPS scores, while donors at the trajectory’s end showed both profoundly altered cellular distributions (Supplementary Figure 7a) and the highest CPS scores (Figure 3f). The scSLIDE trajectory was also associated with additional measures of AD severity including CERAD score (Figure 3g) as well as measures of cognition and dementia (Supplementary Figure 7b,c).

We next investigated cell type abundance shifts along this inferred trajectory and compared our results to binary case-control abundance analyses. We found that specific inhibitory neuronal cell types (such as *SSTe* interneurons) showed clear decreases as AD progressed (Figure 3h, Supplementary Table 3), while among non-neuronal cell types, microglia showed clear upward trends in severe AD patients (Figure 3i). These findings align with established AD pathophysiology, where neuronal loss accompanies reactive gliosis^50,51^. Notably, scSLIDE’s trajectory-based approach captured these changes with strong statistical significance, whereas simple case-control comparisons exhibited trends that did not survive multiple testing corrections (Figure 3h,i).

To validate our observations, we independently applied scSLIDE to the Psych-AD dataset (Figure 4a,b). As in SEA-AD, we fitted a principal curve across the first two diffusion components to represent a trajectory of disease progression (Figure 4b). While independently derived CPS scores were not available for this dataset, our inferred trajectory captured heterogeneity within AD patients (Figure 4c and Supplementary Figure 7d) that correlated with neuropathological assessments of AD severity (including CERAD score and BRAAK category) (Figure 4d, Supplementary Figure 7e). Ordering samples along this trajectory also reproduced the same shifts in cell type abundance that we had observed in SEA-AD (Supplementary Figure 7f,g).

**Figure 4.**
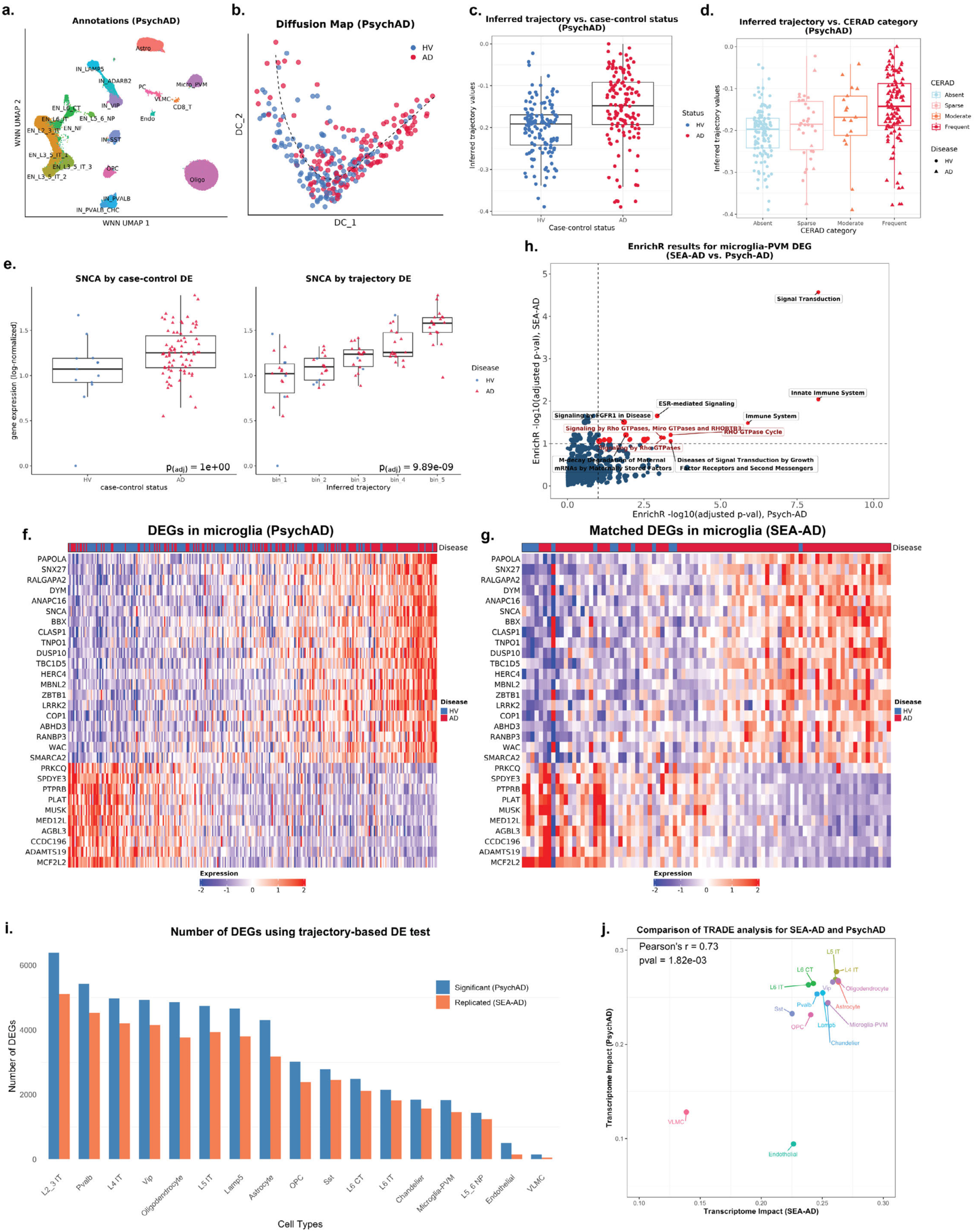
Trajectory-based modeling reveals reproducible molecular programs underlying Alzheimer’s progression. **(a)** Annotated cell-level WNN UMAP of the Psych-AD dataset. Cells are colored based on annotated cell types. **(b)** Diffusion-map embedding and fitted principal curve define a continuous trajectory of disease progression across donors in the Pscyh-AD dataset. **(c,d)** Donor positions along the inferred trajectory relative to case–control status (c) and CERAD plaque burden score (d), showing alignment with pathological severity. **(e)** Example of trajectory-based versus binary case–control differential expression. Expression of SNCA (encoding α-synuclein) shows weak association in case–control testing but strong upregulation along the trajectory (p = 9.89 × 10⁻⁹ with gene-wise Bonferroni correction, see Supplementary Methods). **(f,g)** Heatmaps of trajectory-associated DEGs in microglia identified in Psych-AD (f) and matched DEGs in SEA-AD (g) showing reproducible transcriptional signatures. **(h)** Pathway enrichment of microglial DEGs from SEA-AD (x-axis) and Psych-AD (y-axis), highlighting immune activation and Rho GTPase–mediated signaling linked to disease-associated microglial states. **(i)** Numbers of DEGs per cell type identified by trajectory-based DE analysis in Psych-AD (blue) and nominally replicated in SEA-AD (orange). **(j)** Comparison of transcriptome-wide impact (TI) scores from TRADE analysis across datasets shows strong concordance (Pearson r = 0.73, p = 1.82 × 10⁻³), confirming reproducible cell-type prioritization.

With confidence in the trajectory’s biological relevance across independent cohorts, we next sought to identify the molecular drivers underlying disease progression. We performed trajectory-based DE, modeling each gene’s abundance as a function of donor position along the scSLIDE progression axis (Supplementary Methods). We performed this analysis independently for each dataset, starting with SEA-AD. We found that trajectory based DE analysis was better-suited to model gradual expression changes compared to case-control analysis. As a demonstrative example, we considered *SNCA*, which is a gene encoding α-synuclein that has been causally linked to Alzheimer’s disease in multiple studies^52,53^, with both genetic mutations and microglia-specific overexpression leading to neurodegeneration^54–56^. While we observed non-significant upregulation of *SNCA* in microglia AD patients in binary case-control testing, we identified robust upregulation along the trajectory with high statistical power (Figure 4e).

We further observed the same improvement of statistical power for other genes that have been either genetically or functionally linked to AD, including *SNX27*^57,58^, *LRRK2*^59,60^, *PIK3CA*^61^, *PLAT*^62,63^, and *APOE*^64–66^ (Supplementary Figure 8a, Supplementary Table 4). The increased power also yielded interpretable and biologically meaningful enrichments in microglia, including an enrichment for trajectory-associated genes in both broad immune response and signal transduction pathways (Figure 4f-h). Interestingly, we observed a specific enrichment for genes involved in Rho GTPase pathway activity (Figure 4h). Rho GTPases have been repeatedly linked to aberrant microglial activation and neuroinflammation in Alzheimer’s disease models and have been proposed as therapeutic targets^67–69^, but previous scRNA-seq papers have not identified this enrichment. Thus, the progressive up-regulation of Rho GTPase–related genes along the scSLIDE trajectory represents an interpretable molecular consequence of the transition toward activated, disease-associated cellular states.

Importantly, when we independently performed trajectory-associated DE analysis for the Psych-AD dataset, we observed high reproducibility for each aspect of these expression-based analyses. This includes the reproducible changes in gene expression not just for individual pre-selected AD-associated genes (Figure 4f,g, Supplementary Figure 8a,b), but transcriptome-wide (Figure 4i, Supplementary Figure 9a-c). Interestingly, while we focused our initial analyses on microglia given their likely causal roles in early-stage disease as established via human genetics^70^, we identified substantial transcriptional dysregulation in the majority of cell types (Supplementary Figure 10). As assessed both by the number of DEGs as well as our TI-based prioritization score, we found that only endothelial cells and fibroblast-like Vascular leptomeningeal cells (VLMCs) exhibited minimal transcriptomic changes along the scSLIDE trajectory (Figure 4j). Cell type prioritization was reproducible across independent analyses of both datasets (Pearson’s r = 0.73, p-val = 1.82e-3; Figure 4j), as was the identification of specific DEGs (82.8% reproducibility rate, compared to 17% for case-control DE) across all cell types (Figure 3a, Figure 4i).

Our results strongly suggest that the lack of reproducibility in AD case-control studies arises from the suboptimal treatment of AD as a binary phenotype. Not only is there extensive heterogeneity among cases themselves, but the precise threshold used to separate cases from controls may differ from one study to the next, further limiting comparability. scSLIDE addresses both issues by learning a continuous severity trajectory directly from molecular data.

### Benchmarking scSLIDE performance against alternative approaches

To our knowledge, scSLIDE is the only available method for semi-supervised sample-level embedding. This step is critical for guiding downstream analyses to capture subtle cellular phenotypes that might otherwise be obscured in unsupervised representations. To assess performance and robustness, we assessed and benchmarked scSLIDE across complementary scenarios. First, we performed a negative control analysis to test whether scSLIDE spuriously detects structure when none exists. We randomly permuted sample identities for each cell in the Psych-AD dataset, generated 200 synthetic samples, and randomly assigned case/control labels to each synthetic sample to eliminate true biological differences. We then applied the full scSLIDE workflow, followed by both diffusion map and principal component analysis (PCA) reductions. The resulting cell-level PLS components, sample-level diffusion components and principal components showed no separation of cases and controls (Supplementary Figure 11a-c). A principal curve fit through this structure yielded no differentially expressed genes along the trajectory. This is consistent with the absence of biological signal, indicating that scSLIDE does not overfit (Supplementary Figure 11d).

Next, we compared scSLIDE with five alternative sample-level methods: MRVI^71^, scPoli^72^, PILOT^73^, an unsupervised version of scSLIDE, and a baseline cell-type-proportion workflow (Supplementary Methods). Each approach produces a sample-level distance matrix, which we embedded using diffusion maps^7^. We have previously demonstrated that in AD analysis, scSLIDE identifies a continuous sample trajectory that distinguishes cases and controls, and reproducibly correlates with independent severity metrics. We therefore assessed the ability for each of these alternative methods in recovering these same sources of variation. We note that the SEA-AD dataset^30^ represents a strong test set for benchmarking as the sample-level phenotypes are subtle but representative of complex disease, and an independent ground-truth measure (i.e. CPS score) is available.

When generating a diffusion map based on the sample distance matrix produced by each of the alternative methods, we found that none were capable of reproducing the severity trajectory identified by scSLIDE (Supplementary Figure 12a). To confirm that this was not due to an overt reliance on the first two dimensions, we compared the ability of each method to predict either case-control status or CPS based on the first five diffusion components, using McFadden’s pseudo-r^2^. We found that scSLIDE substantially outperformed all other approaches for both case-control prediction (McFadden’s pseudo r^2^=0.701 for scSLIDE, range of 0.189-0.330 for other approaches), and CPS prediction (r^2^=0.526 for scSLIDE, range of 0.321-0.453 for other approaches; Supplementary Figure 12b,c). Together, these analyses demonstrate that scSLIDE achieves best-in-class performance compared to other methods. Notably, the unsupervised version of scSLIDE failed to recover the same signals, confirming that the supervised dimensional reduction is essential for its improved sensitivity and interpretability for subtle phenotypes such as disease severity. We conclude that scSLIDE delivers state-of-the-art sample-level embeddings that uniquely reveal biologically meaningful heterogeneity across diverse settings.

### Reconstructing developmental heterogeneity within and across embryonic timepoints

Having demonstrated the applicability of scSLIDE across multiple disease contexts, we next turned to a distinct use case of vertebrate development where significant sample heterogeneity is also expected. We examined a recently published snRNA-seq dataset of zebrafish embryogenesis (ZSCAPE)^34^ spanning 18 timepoints (from 18 to 96 hours post fertilization (HPF)), in which major coordinated changes in cellular composition and state occur across all lineages during early development. Importantly, this dataset was collected using the sci-RNA-seq3 technology combined with sci-Plex hashing of individual embryos, where each of ∼528,000 cells was associated with one of 1,025 different wild-type embryos^34^. This design enables us to analyze how sample-level (i.e., embryo-level) heterogeneity unfolds both within individual stages and across developmental time.

We applied scSLIDE to the full zebrafish embryogenesis dataset (Figure 5a, Supplementary Figure 13a), providing the method with discrete timepoint labels but no information about their chronological order (Supplementary Method). Sample-level *t*-distributed stochastic neighbor embedding (*t*-SNE) visualization^74^ and clustering in high-dimensional space revealed that embryos clustered cleanly by developmental stage (Figure 5b,c), with sharper separation than when clustering based only on manually annotated cell-type proportions, as performed in the original manuscript (Supplementary Figure 13b). Notably, scSLIDE also flagged a small number of embryos from HPF 24, 30, 36 as outliers based on their low correlation with other samples from the same timepoint (Figure 5c, Supplementary Figure 13c). Upon closer inspection of sample metadata, we found that these embryos had been cultured at a different starting temperature, a condition known to affect lineage output but not supplied to scSLIDE^75^. We removed these embryos from downstream analyses, illustrating that just as single-cell workflows can detect outlier cells, scSLIDE can detect and exclude outlier samples.

**Figure 5.**
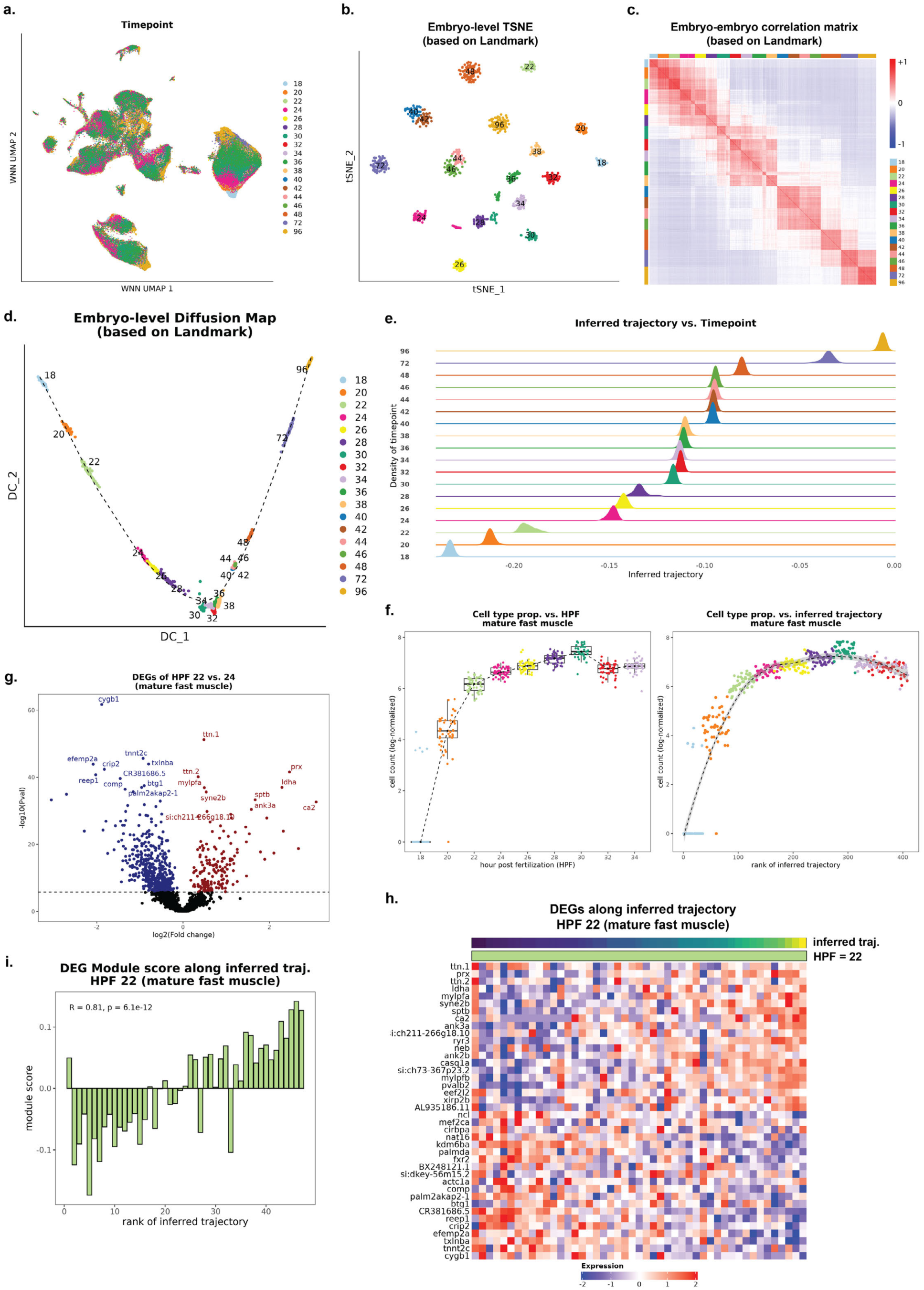
scSLIDE reconstructs developmental heterogeneity during zebrafish embryogenesis. **(a)** Cell-level weighted nearest-neighbor (WNN) UMAP of the zebrafish embryogenesis dataset, with cells colored by developmental timepoint (18–96 hours post-fertilization, HPF). **(b)** Embryo-level t-SNE embedding based on the scSLIDE landmark density matrix reveals sharp separation of developmental stages. **(c)** Embryo-embryo correlation matrix calculated based on the scSLIDE embryo-level relative density matrix. **(d)** Diffusion-map embedding with a fitted principal curve defines a continuous trajectory of developmental progression across embryos. **(e)** Density plots showing the distribution of inferred developmental progression among different embryos, split across timepoints. The inferred developmental trajectory increases with HPF, accurately recapitulating the true temporal order. **(f)** Mature fast-muscle cell abundance across HPF 18–34. (left) shows per-embryo abundance discretely divided by timepoint. (right) orders embryos by their trajectory-inferred progression, revealing gradual transitions in abundance both within and across timepoints. **(g)** Differential expression (DE) between 22 HPF and 24 HPF mature fast muscle identifies known muscle maturation markers. **(h)** Expression of 24 HPF DE genes along the pseudostage trajectory within 22 HPF embryos reveals progressive up-regulation of maturation markers. **(i)** To confirm gradual transcriptional maturation within 22 HPF embryos, we calculated the difference between two module scores along the pseudostage trajectory: the module score for the top 50 up-regulated genes at 24 HPF minus the module score for the top 50 down-regulated genes at 24 HPF.

We next applied diffusion maps to the sample-level relative density matrix to assess whether scSLIDE could capture temporal heterogeneity both within and across timepoints. This revealed a clear developmental trajectory in which embryo progression increased with collection time (Figure 5d,e). We observed nearly identical trajectories when running a fully unsupervised version of scSLIDE that did not use timepoint information (Supplementary Figure 13d), suggesting that in this example (and in contrast to AD), temporal identity was associated with a strong molecular phenotype that did not require supervised analysis.

In addition to capturing heterogeneity across timepoints, scSLIDE also revealed developmental heterogeneity within individual timepoints. As has been described in prior developmental studies^76,77^, such variation can reflect *sample-level pseudostage*, where embryos assigned to the same nominal stage nonetheless occupy distinct positions along the molecular trajectory. Indeed, analogous to single cell pseudo-temporal ordering, we found that sample density exhibited continuous changes along the pseudostage trajectory that were masked in a discrete timepoint-specific view. These changes reflected changes both in cellular proportion across timepoints, but also included variation within a single timepoint that correlated with the pseudostage trajectory (Figure 5f).

For example, mature fast muscle is one of the most dynamic cell types during mid-to-late somitogenesis, with its proportion increasing dramatically between the 18-somite and 24-hour post-fertilization stages as the somitic mesoderm differentiates and fast muscle fibers proliferate^78,79^. When samples were ordered by pseudostage rather than nominal stage, we observed gradual and continuous changes when considering the normalized abundance of pre-annotated cell types for mature fast muscle (Figure 5f). We note that scSLIDE did not leverage cell type annotations or proportions when constructing the trajectory, as it relies on landmark-based estimation of sample density. Similarly, other rapidly changing cell types such as periderm, mature slow muscle, hatching gland, and pharyngeal arch exhibited continuous and gradual changes when aligned by pseudostage (Supplementary Figure 13e-h).

Having established that pseudostage ordering captured within-timepoint variation in cell-type abundance, we next asked whether analogous transcriptomic patterns could be detected. We reasoned that if pseudostage reflects true developmental progression, embryos assigned to adjacent nominal stages should exhibit continuous gene-expression changes. For example, “late” 22-hour embryos should transcriptomically resemble 24-hour embryos. To test this, we compared transcriptomes across consecutive timepoints for matched cell types (Figure 5g–i; Supplementary Methods). In mature fast muscle, genes up-regulated at 24 hours, including *ttn.1*, *ttn.2*, *mylpfa*, and *mylpfb*, known markers of fast-muscle maturation^80–82^, were already elevated in a subset of 22-hour embryos (Figure 5h). Within 22 hpf alone, these genes showed progressive increases along the inferred pseudostage trajectory (Figure 5h,i). This gradual transition was also reflected by corresponding rises in mature fast-muscle cell abundance (Figure 5f). Together, these analyses demonstrate that scSLIDE’s pseudostage recapitulates robust and biologically meaningful sample-level variation even within a single developmental timepoint.

## Discussion

In this study, we introduced scSLIDE, a computational framework aiming to reframe the exploration of heterogeneity in single-cell datasets around the sample rather than the cell. We designed scSLIDE to address key methodological challenges, including defining a cellular embedding that captures both cell type and cell state variation, transforming this into a sample-level representation based on relative high-dimensional density, and leveraging that representation to compute informative sample-level embeddings. Across a variety of basic and translational contexts, we show that scSLIDE enables the grouping of samples based on multiple discrete and continuous sources of variation, the identification of outlier samples, and the characterization of interpretable and reproducible gene expression programs that correlate with sample heterogeneity.

A key advantage of scSLIDE is its ability to resolve multiple, distinct axes of sample-level variation rather than collapsing individuals into binary case-control categories. In our COVID-19 analyses, this framework disentangled independent gradients, such as infection status, time since onset, disease severity that are conflated in traditional workflows. In our analysis of AD, modeling samples along these continuous trajectories substantially boosted statistical power and reproducibility, with transcriptomic signatures replicating across two independent cohorts at rates far exceeding standard case-control methods. More broadly, our results highlight the importance of moving beyond categorical disease labels: many disease processes unfold along gradual and heterogeneous landscapes, and analytical frameworks that explicitly model this structure are essential for accurate and robust inference. By demonstrating that scSLIDE can identify a trajectory that agrees with an independent neuropathology-derived burden score, we hope that scSLIDE will facilitate similar analyses in new contexts.

A distinct aspect of scSLIDE is its use of metadata-assisted supervision in constructing the cell-level embedding. By guiding the embedding toward variation associated with sample-level phenotypes, scSLIDE gains sensitivity to weak or distributed signals that unsupervised methods often miss. Notably, supervision was unnecessary for datasets where temporal processes dominate transcriptional variation, such as zebrafish embryos, yet it proved essential for resolving subtle disease-severity gradients in both COVID-19 and AD. We rely on PLS to implement this supervised component and demonstrate that it remains computationally tractable at atlas scale. However, future extensions could replace or augment this with supervised deep learning approaches, enabling even richer integration of phenotype information as datasets grow in size and complexity.

We also highlight limitations and opportunities for future methodological development. Here, we analyzed multiple large-scale datasets on their own, and while we showed reproducible findings, we did not integrate datasets across studies or consortia. In principle, applying scSLIDE across multi-cohort disease atlases would be extremely powerful, but doing so raises two major challenges. First, while our current workflow mitigates batch effects at the cell level, new approaches will be required to model and correct sample-level batch effects, which may arise from differences in tissue handling, dissociation protocols, or sampling strategies that systematically bias cellular composition. Second, although scSLIDE efficiently handles datasets containing hundreds of samples and more than a million cells, scaling to tens of millions will require further algorithmic advances. Addressing these challenges lies beyond the scope of this study, but represents an exciting direction for future work as sample-rich single-cell datasets continue to expand.

These analyses are made possible by the growing availability of single-cell studies with large numbers of samples. In scRNA-seq, early datasets featuring tens to hundreds of cells could only resolve broad cell classes, but finer-grained states and rare populations only were discovered as cell numbers expanded by orders of magnitude^83^. A similar transition is now occurring at the sample level, particularly with the field’s growing focus on generating datasets with large numbers of patients. As sample sizes continue to grow, we anticipate that sample-level embedding frameworks will reveal increasingly complex patterns including rare outliers, multi-branched trajectories, and clinically relevant subgroups. Moreover, our frameworks can be extended beyond scRNA-seq to single-cell chromatin accessibility, spatial transcriptomics, and multimodal assays, enabling sample-level trajectories to be reconstructed across regulatory and spatial axes. We envision that sample-level embedding frameworks will be essential for translating this next generation of sample-rich single-cell datasets into robust and clinically meaningful insights.

## ACKNOWLEDGEMENTS

We thank all members of the Satija Lab at New York Genome Center for useful discussion. We acknowledge the authors of the external datasets used in this study for making their valuable resources publicly available. This work was supported by the Chan Zuckerberg Initiative (EOSS5-0000000381, HCA-A-1704-01895 to R.S.), and the National Institutes of Health (RM1HG011014-02 and 1OT2OD033760-01 to R.S).

## AUTHOR CONTRIBUTIONS

L.J. and R.S. conceived the research. L.J. performed the computational work and developed the software tool with guidance from R.S and assistance from all authors. All authors participated in interpretation and in writing the manuscript.

## DATA AVAILABILITY

We used publicly available datasets in this work, and the download locations have been described in Supplementary Methods. The COVID-19 COMBAT dataset, Stephenson et al dataset, and SEA-AD dataset are available at CZ CELL×GENE data portal^84^. The Psych-AD dataset can be accessed via AD Knowledge Portal (https://adknowledgeportal.org). The zebrafish embryogenesis dataset can be accessed via the ZSCAPE data portal (https://cole-trapnell-lab.github.io/zscape/).

## CODE AVAILABILITY

Software implementing scSLIDE is freely available as an open-source R package scSLIDE (https://github.com/satijalab/scSLIDE). Vignettes demonstrating the application of scSLIDE are also available as an online resource (https://satijalab.github.io/scSLIDE/index.html).

## SUPPLEMENTARY METHODS

### 1. scSLIDE workflow

scSLIDE is a sample-centric framework for analyzing multi-sample single-cell data. It represents each sample as a distribution over cellular states in a common latent space, transforming a single cell gene expression matrix into a sample-level relative density matrix for downstream clustering, trajectory inference, and differential analyses. As described below, the workflow first constructs a cell-level embedding and subsequently transforms this into a sample-level representation. All methods are implemented in the open-source R package scSLIDE (www.github.com/satijalab/scSLIDE).

#### 1.1 Type-oriented cell embedding

We first obtain a low-dimensional representation aiming to capture high-resolution cell types. scSLIDE is agnostic to the specific approach, as users may supply a PCA-based embedding or an embedding from common integration methods (e.g., “anchor-based” integration^37^, Harmony^38^, scVI tools^39^). Cell type annotations are not utilized by scSLIDE itself, but can be used later to interpret downstream results. As an alternative to unsupervised integration procedures, we recommend the use of reference mapping workflow when a high-quality reference dataset is available. These workflows embed cells in an informative high-dimensional space alongside the reference dataset. scSLIDE supports reference-mapped embeddings produced by methods such as Azimuth^10^ or similar tools like scArches or Symphony^11,40^. By default, our implementation leverages the Pan-Human Azimuth reference (https://satijalab.org/pan_human_azimuth/) developed as part of the NIH Human Biomolecular Atlas Program (HuBMAP) for human sc/snRNA-seq data^85^. Pan-Human Azimuth provides a consistent hierarchical ontology and embeds cells into a 128-dimensional space that aims to preserve high-resolution cell types.

#### 1.2 Phenotype-associated cell embedding via partial least squares (PLS)

Type-focused embeddings preserve cell identity but often under-weight subtle, phenotype-associated state differences (e.g., disease status). scSLIDE adopts a supervised embedding using partial least squares (PLS), which learns components that maximize covariance between the gene expression matrix and sample-level phenotypes repeated at the cell level. Specifically, we adopt the canonical PLS (CPLS) implementation of PLS from ref.^86^, as it has been observed to achieve stronger predictive performance with fewer components for categorical or multi-output phenotypes than standard PLS.

We define the following:

**X**: the log-normalized gene expression matrix with n cells and p features. Each column (feature) has been standardized with mean = 0 and standard deviation = 1.

**Y**: the phenotype matrix with n cells and q outcomes (i.e., sample-level phenotypes repeated across cells from the same donor).

CPLS process can be summarized by the following equation:

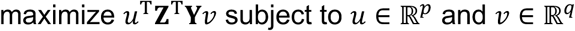

where **Z** = **XX**^T^**Y**. The actual solution is given by:

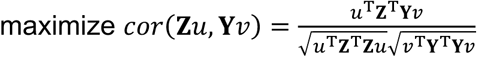

Supervised dimensional reduction is performed by the scSLIDE::RunPLS(…, method = c(”cppls”)) function.

When choosing features (i.e. genes) to use for supervised dimensional reduction tasks, we included the top 2,000 highly variable genes as chosen by default in Seurat. In addition to selecting highly variable genes, we enriched the feature set for PLS by incorporating genes with potential disease relevance. Within each cell type, we performed a simple Pearson correlation test between gene expression and the disease label, ranking genes by the strength of their association. We then selected the top ∼1% of genes per cell type and added them to the variable-gene list. This step ensures that the supervised dimensionality reduction has access to markers that capture phenotype-associated variation even if these genes were not independently identified as highly variable in the unsupervised analysis. We utilized this process during feature selection for all analyses in this manuscript.

#### 1.3 Landmark selection

We define landmarks as a subset of cells that span the full set of cell types and states in the dataset (inspired by scLKME^35^). In scSLIDE, landmarks are the reference points used to quantify how each sample’s cells distribute across cellular neighborhoods (Section 1.4). Landmarks are selected once from the full cell set and then carried through both embeddings (type-oriented and PLS), since each landmark cell has coordinates in both spaces.

As landmarks are meant to describe the distribution of the full dataset, it is essential that they preserve the full transcriptional diversity of the data, including both abundant and rare states. To achieve this, we select landmarks using leverage-score–based sketching, a principled approach for identifying the cells that have the greatest influence on the global gene–gene covariance structure. Intuitively, cells with high leverage scores correspond to transcriptional profiles that define important directions of variation in the dataset, making them valuable representatives for downstream density estimation. In ref.^42^, we showed that selecting cells via this geometric sketching-based procedure successfully identifies cells that encompass the full diversity of rare and abundant cell types and states in a variety of datasets.

We approximate these leverage scores using the Clarkson–Woodruff randomized sketching framework, which provides an efficient alternative to computing exact leverage scores via full matrix decompositions. Briefly (and as fully described in ref.^42^) , we apply a sparse CountSketch projection to the cell-by-gene matrix to reduce dimensionality while preserving key variance components. We then compute a QR decomposition of this sketched matrix and apply a fast Johnson–Lindenstrauss projection to obtain a low-rank embedding from which approximate row-wise leverage scores can be computed as squared Euclidean norms. These scores quantify the contribution of each cell to the global covariance structure, and we sample cells with probability proportional to their leverage score. This procedure enriches the landmark set for rare, structurally informative, and transcriptionally distinct states, while still maintaining broad coverage of abundant cell populations.

For each analysis in the manuscript, we used 5,000 landmarks across the dataset. We showed this is a setting that balances coverage of rare states with runtime and memory. Moreover, we show that scSLIDE’s performance remains robust across a range of landmark numbers (Supplementary Figure 1b).

#### 1.4 Joint embedding with weighted nearest neighbors (WNN)

The type-oriented embedding and phenotype-associated embedding are two embeddings that capture two complementary representations of a single-cell data. Next, we aimed to efficiently combine the two sets of information together, and quantitatively measure the cell density near each landmark for every sample on this joint representation space. We adopted the weighted nearest neighbor (WNN), which we initially developed to integrate multiple modalities measured within a cell and to obtain a joint definition of cellular state^10^. Here, we treat the two embeddings as two different modalities from the same group of cells. We then quantify the cell-landmark distance using a weighted average of their distances on both embedding spaces. We define the following:

*c_type_*: L2 normalized low-dimensional vector representing the type-oriented embedding for a cell.

*l_type_*: L2 normalized low-dimensional vector representing the type-oriented embedding for a landmark

*c_p_*_,-_: L2 normalized low-dimensional vector representing the PLS embedding for a cell

*l_pls_*: L2 normalized low-dimensional vector representing the PLS embedding for a landmark

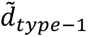: the Euclidean distance between a cell and its first nearest landmark (i.e., the local connectivity) on the type-oriented embedding

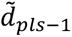: the Euclidean distance between a cell and its first nearest landmark (i.e., the local connectivity) on the PLS embedding

*σ_type-1_*: the Euclidean distance between a cell and its k-th nearest (k = 20 by default) landmark on the type-oriented embedding; used as the cell-specific kernel bandwidth

*σ_type-k_*: the Euclidean distance between a cell and its k-th nearest (k = 20 by default) landmark on the PLS embedding; used as the cell-specific kernel bandwidth

The WNN procedure can be described as follows:

i. Build k-NN graphs (k = 200) from cells to landmarks separately for the type-oriented embedding and phenotype-associated embedding, respectively. The k-NN is based on euclidean distances (with L2 normalization) between cells and landmarks:

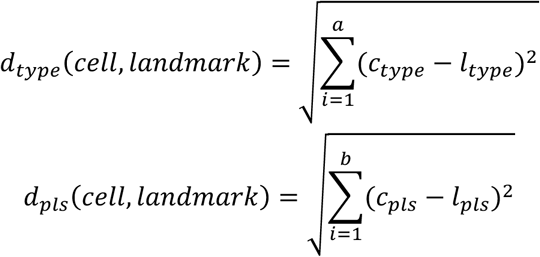
ii. Convert distances to affinities with an exponential kernel and the standard local-connectivity correction^87^:

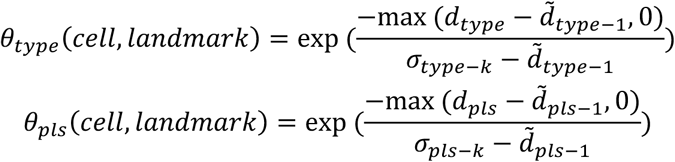
iii. Combine the two affinities into a WNN affinity by a fixed average:

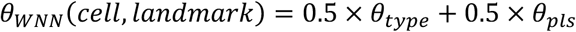

We choose to set the two embedding weights to 0.5 and 0.5 to balance the information from both embeddings in downstream analyses, and use these default values for all analyses in this manuscript.

From these affinities, scSLIDE builds a WNN graph linking each cell to its top-k landmark neighbors (default k=5). For each cell, we consider the set:

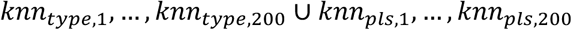

and identify the top k (k = 5 by default) landmark WNN neighbours.

The process of WNN graph construction is implemented in the scSLIDE::PrepareSampleObject(), which returns a landmark-by-cell WNN graph.

#### 1.5 Sample-level density matrix construction and chi-square normalization

From the asymmetric landmark-by-cell WNN graph, scSLIDE constructs a landmark-by-sample density matrix by first converting each landmark–cell connection (from the previously generated WNN graph) into a binary indicator, assigning a value of 1 when a landmark is identified as falling within the *k-*nearest neighbors (*k*=5 by default) of a cell and 0 otherwise. Each cell is associated with a specific sample, and scSLIDE aggregates these binary indicators by summing them across all cells belonging to the same sample. In effect, this procedure counts how many cells from each sample lie in the local high-dimensional neighborhood of each landmark, producing a matrix (**L**) in which each entry reflects the density of a sample’s cells near a given cellular state. This matrix serves as a compact, sample-level representation of how each donor’s cells are distributed across the full landscape of cellular types and states captured by the landmarks.

The values in the resulting matrix vary systematically with each sample’s total number of profiled cells and therefore are not directly comparable. To correct for these library-size differences and harmonize the representations across samples, we normalize the matrix using a chi-square–based residual transformation. The transformation adjusts each entry by comparing the observed count to the expectation under a model with no biological differences, yielding values that reflect true enrichment or depletion near each landmark rather than variation in sampling depth.

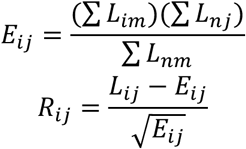

Note that **E**m_×_n is the expected chi-square table under the null (i.e., no relationship between any two samples). The resulting harmonized table **R**m_×_n is the **sample-level relative density matrix**. Positive residuals indicate that a sample is enriched for cells near a given landmark, while negative residuals indicate depletion. **R**m_×_n can be viewed as a normalized donor profile that supports standard downstream analyses at the sample level, including distance-based similarity, PCA, clustering of samples, trajectory inference (diffusion map analysis), etc. This step is implemented in our scSLIDE::GenerateSampleObject() function.

#### 1.6 Achieving computational scalability on large datasets

A central goal of scSLIDE is to enable sample-level analyses on datasets that include hundreds of donors and millions of cells, a scale at which both memory usage and computational cost can become prohibitive. To accommodate these demands, we implemented several strategies that ensure computational stability and efficiency for scSLIDE.

First, scSLIDE is fully compatible with the BPCells–Seurat v5 infrastructure^42,88^, which enables out-of-memory storage of massive single cell datasets. Using this framework, large scRNA-seq datasets can be stored on disk, and only the necessary subsets of cells or features are loaded into memory at any given time. This design allows scSLIDE to process datasets containing millions of cells on standard computational hardware.

The most computationally intensive step of the workflow is the PLS-based supervised dimensionality reduction. PLS requires repeated computation of covariance structures between the gene expression matrix and the phenotype matrix, which can be expensive when applied to all cells. To improve efficiency, we adopt a two-step strategy. For each sample, we randomly select 3,000–4,000 cells and train the PLS model on this subset. Once the PLS components are obtained, all remaining cells (those not included in the random subsample) are projected onto the learned PLS loadings. This dramatically reduces runtime while preserving the key phenotype-associated variation needed for downstream analyses. We also found that this subsampling strategy mitigates potential imbalances in cell numbers across donors and yields more stable phenotype-associated components. Although technically optional, this procedure is used for all analyses in this manuscript and is set as the default behavior in scSLIDE.

Together, these strategies result in a highly scalable workflow. As a representative benchmark, scSLIDE processed the Psych-AD dataset (∼300 donors and ∼1.2 million cells) in approximately 2 hours (with 4 CPU cores), including all steps from embedding construction to sample-level density estimation. These optimizations position scSLIDE to scale with the increasing size of modern single-cell datasets while maintaining robust and efficient performance.

### 2. Sample-level diffusion maps and pseudotrajectory inference

The sample-level relative density matrix provides a normalized profile for each donor and supports downstream analyses. While PCA can reveal major axes and enable clustering, it is a linear decomposition, and may not perform well when variation follows curved or branching manifolds (common in temporal or severity gradients)^7,89,90^. Diffusion maps address this by constructing a Markov process over the data graph, emphasizing connectivity along the manifold and yielding coordinates (diffusion components, DCs) that capture continuous progressions. While typically the diffusion map is utilized to infer pseudo-trajectories in single-cell data, here we applied it to our sample-by-landmark relative density matrix to arrange the “samples” along a pseudo-trajectory based on their pairwise similarity. In practice, we implemented the destiny::DiffusionMap() function from the “destiny” R-package^91^ to perform the diffusion map analysis. The distance metric is set to “cosine”, and the distance is calculated using all the available landmarks. The final DCs are weighted by the root of the corresponding eigenvalues. All of these have been implemented in the scSLIDE::RunDiffusionMap() wrapper function.

### 3. Trajectory-based differential expression (DE) analysis

After calculating a sample ordering (for example, based on their position along a diffusion component), we test for genes whose expression changes along this continuous axis. We reasoned that treating progression as a continuous predictor should increase power compared to binary case-control testing when disease effects are gradual, heterogeneous across donors, or only partially aligned with the case/control label. We therefore developed a negative binomial generalized linear model (NB-GLM) using donor-level pseudobulk counts as the response and diffusion component (or trajectory) scores as the predictor. Sample-level pseudobulk counts are generated for each cell type separately using Seurat::AggregateExpression(). As for the trajectory DE test based on NB-GLM, we define the following:

*y*: vector of donor-level pseudo-bulked RNA-seq expression counts for a response gene. The pseudo-bulked counts can be obtained by the Seurat::AggregateExpression() function in a donor- and cell type-specific manner (i.e., the donor-level pseudo-bulking is performed for each cell type individually).

*s*: vector of the trajectory scores.

*Z*: matrix of covariates (including an intercept term). In our study, covariates include age, sex, and log_10_(total counts).

And we fit the following negative binomial regression model:

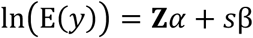

where *y* ∼ NB(*μ*, *θ*) and 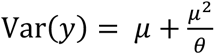 . Our parameter of interest is β, the coefficient for the progression score. A likelihood-ratio test is used to evaluate the null hypothesis H_9_: β = 0.

For implementations, model fitting and likelihood-ratio test is performed using the glmGamPoi::glm_gp() function from the glmGamPoi R package (v.1.10.2)^92^. The wrapper scSLIDE::TrajDETest() takes the pseudobulk counts, the trajectory vector, and relevant covariates and returns per-gene estimates and statistics. To offer an interpretable metric of the magnitude of the effect, we report the log-fold change of each gene, which is computed between donors in the top vs. bottom tertiles of *s* (default: upper and lower 1/3). P-values are adjusted gene-wise and by default we use a Bonferroni threshold throughout the paper (e.g., 0.05 / 30,000). scSLIDE also returns FDR (Benjamini–Hochberg) for users who prefer FDR control in exploratory analyses.

### 4. Differential abundance (DA) analysis

We performed differential abundance analysis using the inferred diffusion component or pseudotrajectory. We first quantified the number of cells for each cell type in each donor to obtain a count matrix (C), then performed log-normalization for each sample:

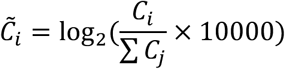

This normalization accounts for varying total cell counts across samples and converts raw counts into log-normalized proportions (following recommendations from ref.^93^). For each cell type, we performed a Pearson correlation test between cell type proportions and trajectory scores. Cell types were ranked by correlation coefficients and p-values, and top positively and negatively correlated cell types were reported after Bonferroni correction (threshold: 0.05/number_of_cell_types).

### 5. Transcriptome-wide impact (TI) quantification by cell type

Our trajectory DE test prioritizes cell-type-specific DE genes relevant to disease trajectories but cannot quantitatively summarize the overall transcriptome shift for each cell type. To identify which cell types are most transcriptionally affected, we adopted TRADE^46^ to evaluate the transcriptome-wide impact (TI) of each diffusion component (or pseudotrajectory) on each cell type. TRADE is a mixture model that uses summary-level effect sizes (log2 fold changes) and standard errors from case-control DE tests to estimate the distribution of true differential expression effects^46^. It does not rely on arbitrary significance thresholds but instead estimates the distribution of true DE effects using all genes and summarizes it into a single TI value (the variance of the effect size distribution) representing the overall transcriptome shift.

TRADE assumes effect sizes from standard case-control DE tests (cases coded as 1, controls as 0), which are naturally in log2 fold change units^94^. To adapt TRADE to our framework, we adopted a standardization method common in human genetics^95^, using DE test z-scores and sample sizes to re-scale the effect sizes and standard errors. Specifically, after obtaining DE test summary results, we performed the following conversion:

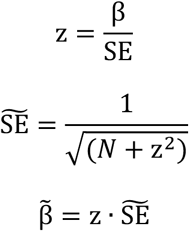

where β and SE are the original effect size and standard error, and N is the sample size. β̃ represents the effect size when gene expression is in standard deviation units, and 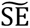 is the standardized SE. By applying this conversion to all genes, their β̃ share the same scale and can be used for TRADE analysis. For each cell type, we performed this standardization followed by TRADE analysis to obtain TI values for prioritizing cell type relevance to the inferred trajectory.

We also performed Augur^47^ analysis as an alternative cell type prioritization method. Augur is a machine learning framework that quantifies the separability of perturbed and unperturbed cells using single-cell data and case-control labels. Following default guidelines, we provided cell-type-specific single-cell data and case-control labels to obtain an AUC-based prioritization score for each cell type.

### 6. Pathway enrichment analysis

To interpret DEGs from our DE test, we performed pathway enrichment analysis using the EnrichR R-package^96^ with default settings. Enrichment analysis identifies biological processes and pathways overrepresented in a gene list. For each cell type, we provided statistically significant DEGs (after gene-wise Bonferroni correction) as the target gene list and all expressed genes in that cell type as the background^97^. Enrichment was performed against the Reactome Pathway 2024 database^98^.

### 7. Applications to real datasets

#### 7.1 COVID-19 COMBAT (Ahern et al) dataset

We downloaded the COVID-19 multi-omics blood atlas (COMBAT) scRNA-seq dataset^14^ in .h5ad format from the Chan Zuckerberg Cell×Gene data portal^84^. We annotated the dataset using the Pan-Human Azimuth python package (https://satijalab.org/pan_human_azimuth/) which provided cell type labels at multiple granularities and 128-dimensional cell-level embeddings. We then used the on-disk functionality of Seurat v5^42^ and BPCells^88^ to load the .h5ad file into R, generating an on-disk Seurat object with appended metadata and Azimuth embeddings.

We performed quality control and removed cells with <500 detected genes, <1,000 UMI counts, or cells that failed to pass Azimuth QC (inconsistent cell type hierarchy or low prediction confidence <0.6). We retained cells from healthy controls (”HV”, n=10 donors) and COVID-19 patients (”COVID_MILD”: mild, n=18; “COVID_SEV”: severe, n=41; “COVID_CRIT”: critical, n=18), excluding other disease categories. Any individuals with <200 total single cells were removed. The final QC-filtered dataset contained 499,027 cells. For supervised dimensional reduction (PLS), the disease phenotypes (”HV”, “COVID_MILD”, “COVID_SEV”, “COVID_CRIT”) were coded as dummy variables (Y matrix).

#### 7.2 COVID-19 Stephenson et al pNewcastlep dataset

We downloaded the COVID-19 Stephenson et al. “Newcastle” scRNA-seq dataset^33^ in .h5ad format from the Chan Zuckerberg Cell×Gene data portal^84^. As with the Ahern dataset described above, we annotated the data using the Pan-Human Azimuth python package and generated an on-disk Seurat object. We performed QC to remove cells with <500 detected genes and <1,000 UMI counts. The Stephenson et al. “Newcastle” dataset used a different COVID-19 severity classification: “Healthy” (n=12 donors), “Asymptomatic” (n=1), “Mild” (n=11), “Moderate” (n=17), “Severe” (n=7), and “Critical” (n=7). We retained these original severity categories and utilized them for the PLS dimensional reduction. The final QC-filtered dataset contained 366,357 cells.

#### 7.3 Alzheimer’s disease SEA-AD dataset

We downloaded the Alzheimer’s disease SEA-AD single-nucleus RNA-seq (snRNA-seq) dataset^30^ in .h5ad format from the Chan Zuckerberg Cell×Gene data portal^84^. We used high-confidence cell type labels provided by the original study^30^ and obtained cell-level Azimuth embeddings using the Pan-Human Azimuth python package. We generated an on-disk Seurat object and performed QC to remove nuclei with <500 detected genes, <1,000 UMI counts, or cells that failed to pass Azimuth QC (inconsistent cell type hierarchy or low prediction confidence <0.6). Individuals with <200 nuclei were removed. The remaining dataset consists of donors that were labeled as “healthy” (n=14) or “AD” (n=75). The final QC-filtered dataset contained 1,309,968 nuclei.

#### 7.4 Alzheimer’s disease Psych-AD dataset

We downloaded the Alzheimer’s disease Psych-AD snRNA-seq dataset^25^ in .h5ad format from the AD Knowledge Portal (https://adknowledgeportal.org) and Synapse data portal (www.synapse.org), project ID syn51188606. We used high-confidence cell type labels from the original study and obtained cell-level Azimuth embeddings using the Pan-Human Azimuth python package. We generated an on-disk Seurat object and performed QC to remove nuclei with <500 detected genes and <1,000 UMI counts. Donors were labeled as “healthy” (n=147) or “AD” (n=145). The final QC-filtered dataset contained 1,219,003 nuclei.

To evaluate cross-dataset consistency between SEA-AD and Psych-AD, we compared DE test z-scores for each cell type. We also compared DA results and TRADE rankings between datasets. For standard case-control DE testing, we used Psych-AD as the discovery dataset, extracted significant DEGs after gene-wise Bonferroni correction, and quantified the proportion of nominally replicated DEGs (raw p-value ≤0.01) in SEA-AD. We performed the same replication analysis for trajectory-based DE results.

#### 7.5 ZSCAPE zebrafish embryogenesis dataset

We downloaded the ZSCAPE zebrafish embryogenesis single-nucleus sci-RNA-seq3 dataset^34^ in .mtx format from the ZSCAPE data portal (https://cole-trapnell-lab.github.io/zscape/). We obtained the reference atlas containing wild-type controls, loaded the .mtx files into Python to generate an AnnData^99^ object, and saved it in .h5ad format. We generated an on-disk Seurat object and performed QC to remove nuclei with <100 detected genes or <200 UMI counts. Embryos with <200 nuclei were removed. We retained nuclei from uninjected control embryos and excluded other experimental conditions. The final QC-filtered dataset contained 528,695 nuclei.

To quantify developmental heterogeneity within individual timepoints, we performed pairwise comparisons between adjacent timepoints. For each cell type, we conducted standard pseudobulk case-control DE testing (earlier timepoint coded as 0, later timepoint as 1) to identify significant DEGs. Within the earlier timepoint, we ordered embryos by their inferred scSLIDE trajectory and quantified: (1) expression levels of up- and down-regulated case-control DEGs, (2) module scores^100^ for the top 50 up- and down-regulated DEGs, and (3) cell type proportions. For comparison, we also analyzed the data using a standard cell type proportion based strategy as motivated by the original study (see section 9)^34^. Briefly, we calculated the number of cells per cell type for each embryo, generating a cell-type-by-embryo count matrix, and applied log-normalization (Seurat::NormalizeData() with default parameters, following recommendations from ref.^93^) to obtain log-transformed proportions for downstream analyses.

### 8. Null simulations

To assess whether scSLIDE is well-calibrated under the null hypothesis of no biological signal, we performed null simulations using healthy individuals from the Psych-AD dataset. We randomly permuted donor identities across nuclei to create 200 synthetic donors and randomly assigned case/control labels (1:1 ratio) to each. We then applied the full scSLIDE workflow. For the synthetic sample-level relative density matrix, we performed both diffusion map analysis and principal component analysis (PCA). We fitted a principal curve to the top two PCs to construct a pseudotrajectory for trajectory-based DE testing. We generated quantile-quantile (Q-Q) plots of DE test p-values for each cell type to assess calibration under the null hypothesis (under proper calibration, p-values should follow a uniform distribution).

### 9. Benchmarking analysis

To assess scSLIDE’s performance, we compared it with recently proposed sample-level analytical frameworks: MRVI^71^, scPoli^72^, PILOT^73^. We also included two control methods: an unsupervised version of scSLIDE (u-scSLIDE, which constructs cell-by-landmark k-nearest neighbor graphs (k=5) using only the Pan-Human Azimuth embedding without PLS-derived information) and a cell-type-proportion-based workflow. All methods generate either sample-level distance matrices (MRVI, PILOT) or sample-level embedding matrices (scPoli, scSLIDE, u-scSLIDE, cell-type-proportions), which can be converted to distance matrices. We applied diffusion map analysis to each method’s sample-level matrix using scSLIDE::RunDiffusionMap() to obtain diffusion components (DCs) for downstream comparisons.

We benchmarked methods on the Alzheimer’s disease SEA-AD dataset. SEA-AD contains >1.3 million nuclei after QC and exceeds computational limits for some methods, so we randomly subsampled 50% of nuclei and applied all methods (including scSLIDE for fair comparison) to this subset.

After obtaining the sample-level distance matrix (or sample-level embedding matrix) from each method, we used it as input for diffusion map analysis and extracted the top five diffusion components (DCs) for downstream benchmarking. For SEA-AD, we fitted logistic regression models using the top five DCs as predictors and case-control status as the outcome, reporting McFadden’s pseudo-r² to compare predictive performance across methods. We additionally fitted linear regression models using the top five DCs to predict the continuous pseudo-progression scores (CPSs) provided by the original study and reported the standard r².

For all external methods, we followed recommended settings. Detailed implementations are described below.

#### 9.1 MRVI

MRVI is an unsupervised deep generative model that generates sample-sample distance matrices from scRNA-seq data for exploratory analysis of sample stratifications^71^. We used scvi-tools v1.3.1^101^ with default settings (https://scvi-tools.readthedocs.io/en/latest/index.html). Following the tutorial, we selected the top 10,000 highly variable genes for model training. After training, we computed sample-sample distance matrices for each cell type separately and for all cells pooled together. Distance matrices were imported into R for diffusion map analysis.

#### 9.2 scPoli

scPoli is a conditional variational autoencoder (CVAE)-based model that learns both cell-level and sample-level representations from single-cell data^72^. Although designed primarily for data integration, scPoli can generate sample-level embeddings by setting the “batch” parameter to sample identity. We used scArches v0.6.1^11^ with default settings (https://github.com/theislab/scarches). Following the tutorial, we selected the top 4,000 highly variable genes for training. After training, we extracted the sample-level embedding matrix and imported it into R for diffusion map analysis.

#### 9.3 PILOT

PILOT is an optimal transport-based method that models each sample’s single-cell data as a distribution in gene expression space and computes sample-sample Wasserstein distances^73^. We used PILOT v2.0.6 with default settings (https://pilot.readthedocs.io/en/latest/index.html). We selected the top 4,000 highly variable genes and performed PCA to obtain the top 50 principal components. PILOT used these 50 PCs to define cell state clusters and represented each sample as a distribution over these clusters. Sample-sample Wasserstein distances were computed using optimal transport with costs based on cluster similarity in PC space.

#### 9.4 Unsupervised scSLIDE (u-scSLIDE)

u-scSLIDE is an unsupervised variant of scSLIDE that omits PLS training and uses only Azimuth embeddings for sample-level representation, serving as a control to assess the contribution of supervised dimensionality reduction. After Pan-Human Azimuth annotation, QC, and preprocessing, we performed geometric sketching to select 5,000 landmark cells. We then constructed a landmark-by-cell k-nearest neighbor graph (k=5) using only the Azimuth embedding (128 dimensions) via Seurat::FindNeighbor(). The kNN graph was aggregated into a landmark-by-sample density matrix, which was then chi-square normalized to obtain relative densities for downstream diffusion map analysis.

#### 9.5 Cell-type-proportion-based workflow

We included a cell-type-proportion-based workflow as a simple baseline that captures only cell type composition changes. After QC, we calculated the number of cells per cell type for each donor, generating a cell-type-by-donor count matrix. We then applied log-normalization (Seurat::NormalizeData() with default parameters, following recommendations from ref.^93^) to obtain log-transformed proportions for diffusion map analysis.

