## Supplemental Figures for "Reconstructing developmental and disease progression with sample-level embeddings"

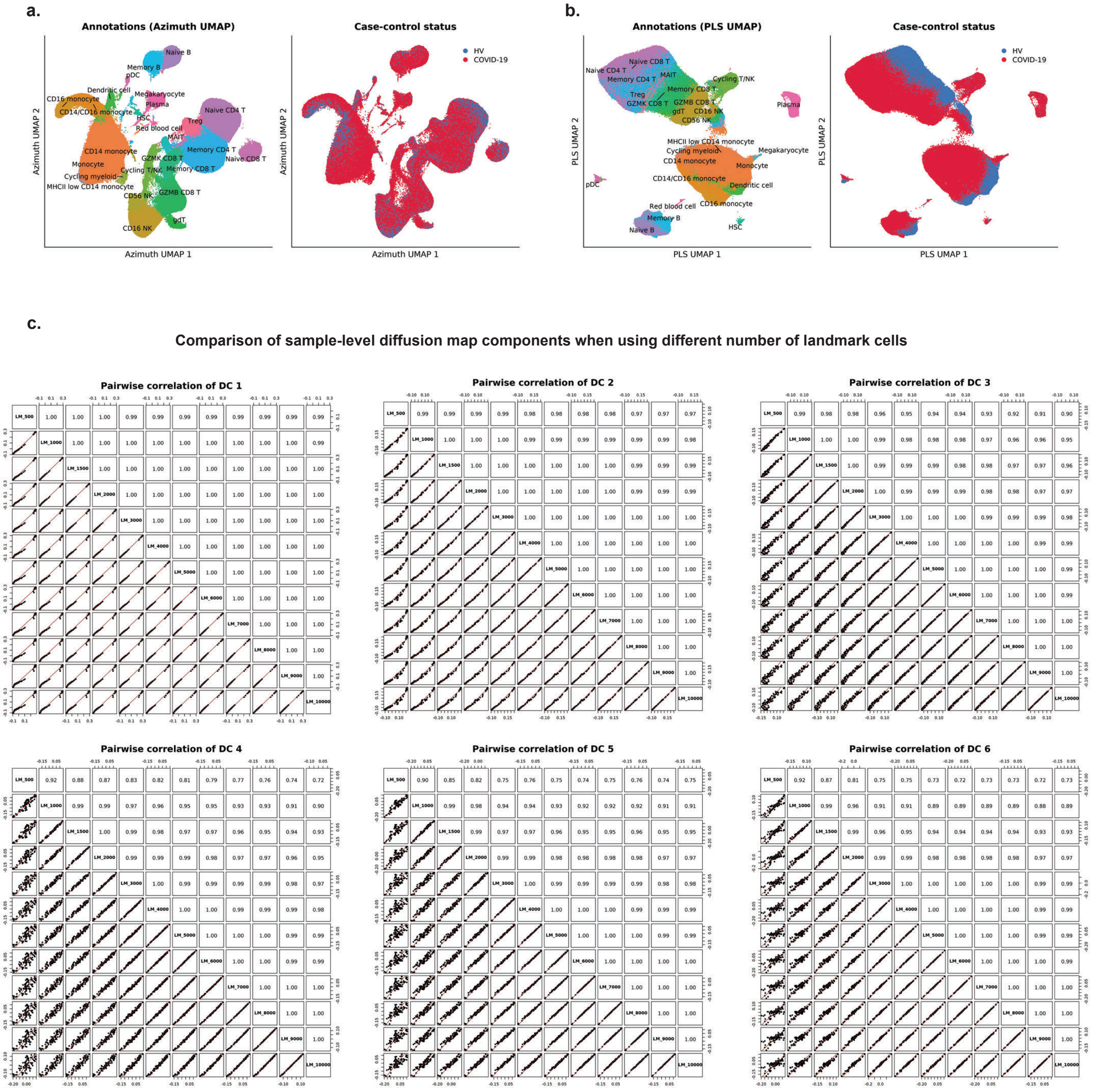


**Supplementary Figure 1.** (a) UMAP of the *cell-type–oriented* embedding obtained by mapping COMBAT COVID-19 cells to the Azimuth reference, illustrating that the embedding preserves fine-grained immune cell structure. (b) UMAP of the *disease-associated* embedding derived using partial least squares (PLS), highlighting variation linked to COVID-19 infection and disease response. (c) Comparison of sample-level diffusion map components across a wide range of landmark set sizes (500–10,000 cells). The high concordance across settings shows that scSLIDE is robust to the choice of the exact number of landmarks. We utilize 5,000 landmarks for each analysis in this manuscript.

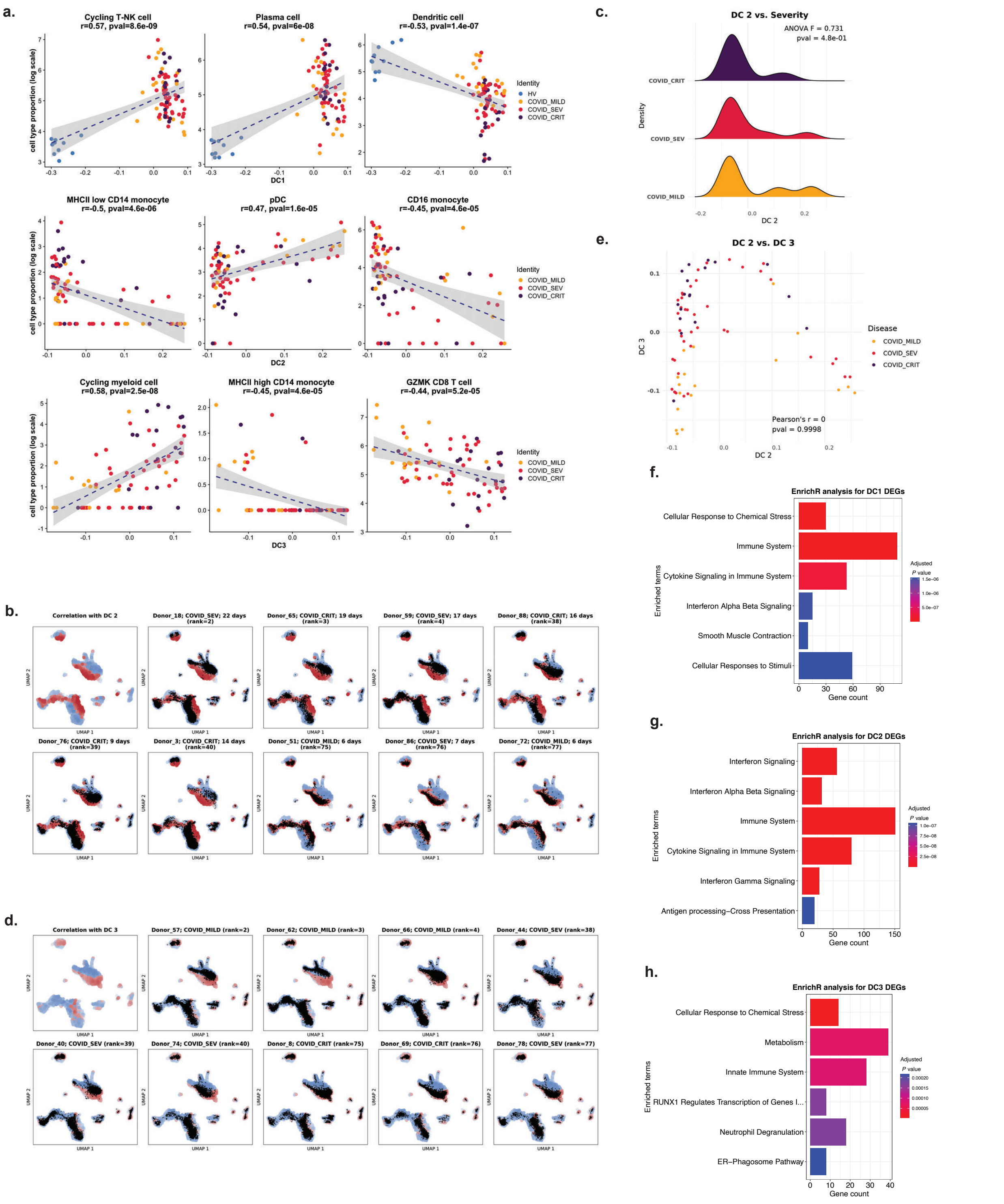


**Supplementary Figure 2.** (a). Changes in cell-type proportions across the different diffusion components (DCs) of the sample-level density matrix for the COMBAT COVID-19 dataset. (b) Visualizing the density of cells from each individual sample. Each colored point represents a landmark, and is colored by Pearson correlation (red, positive; blue, negative) with DC2. (c). A ridge plot showing DC2 does not correlate with COVID-19 severity. (d). Similar to (b) but showing the density of cells from each individual along DC3. (e). Diffusion map embedding of the sample-level relative density matrix for DC2 and DC3, showing that the DC2 axis is independent from the DC3 axis. (f-h). EnrichR pathway analyses for DC-associated differentially expressed genes (DEGs): DC3 emphasizes neutrophil degranulation and related inflammatory programs; DC2 is dominated by interferon-stimulated gene signatures and antiviral pathways; DC1 reflects broad immune/inflammatory responses and cellular stress programs. The gene ontology databases being used is REACTOME (10.1093/nar/gkad1025).

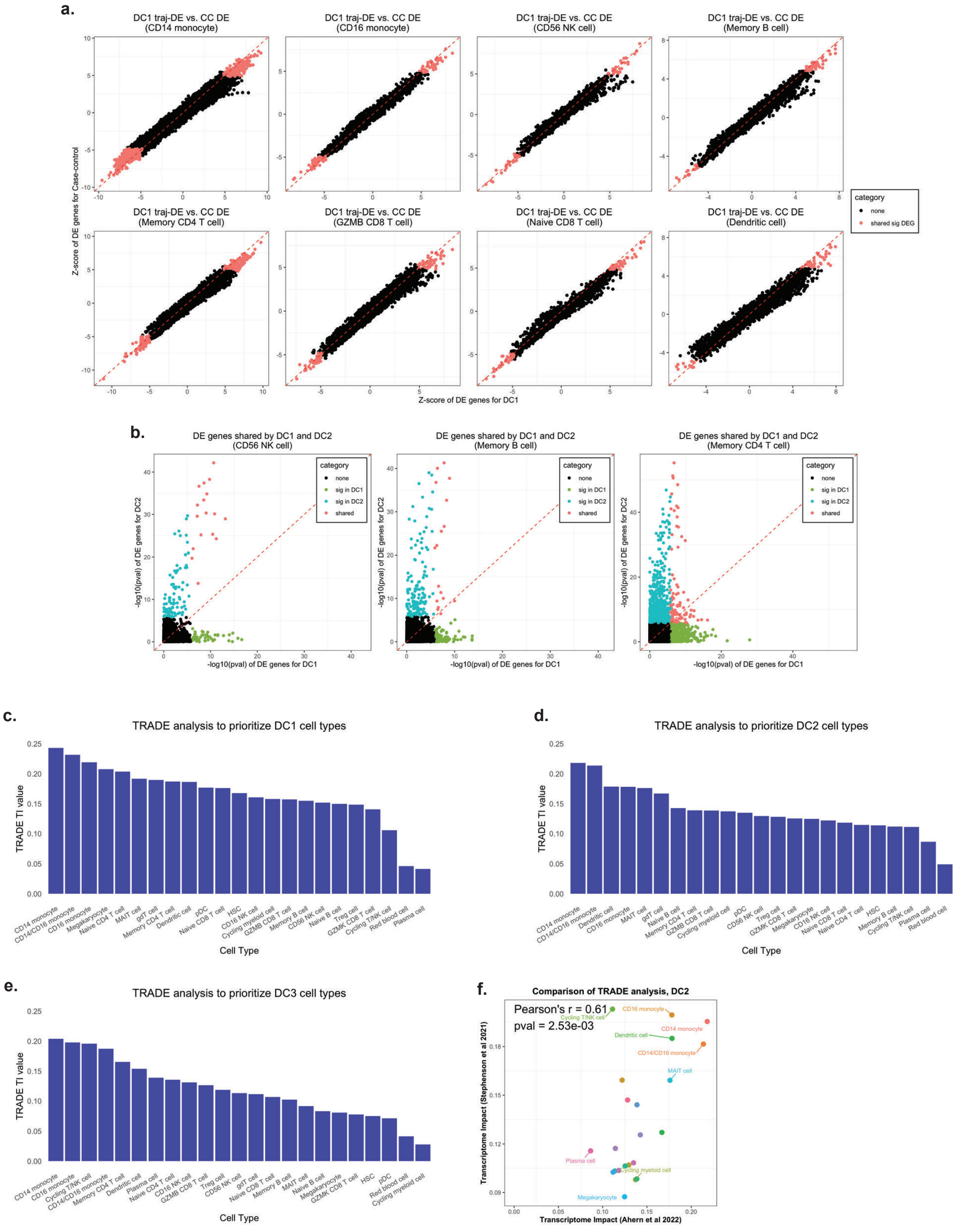


**Supplementary Figure 3.** (a) Scatter plots showing consistency of DE test z-scores between a standard pseudo-bulked case-control DE test and a trajectory-based pseudo-bulked DE test for DC1 for the COMBAT COVID-19 dataset. (b) Minimal overlap between DC1/2 DE gene sets indicates that each component captures distinct biological axes of variation. (c–e) Transcriptome Impact (TI) scores from TRADE analysis (Supplementary Methods) across cell types for DC1–DC3, for the COMBAT dataset. (f) Comparison of transcriptome-wide impact (TI) scores for DC2 from TRADE analysis across cell types shows strong reproducibility between the independent Ahern and Stephenson datasets.

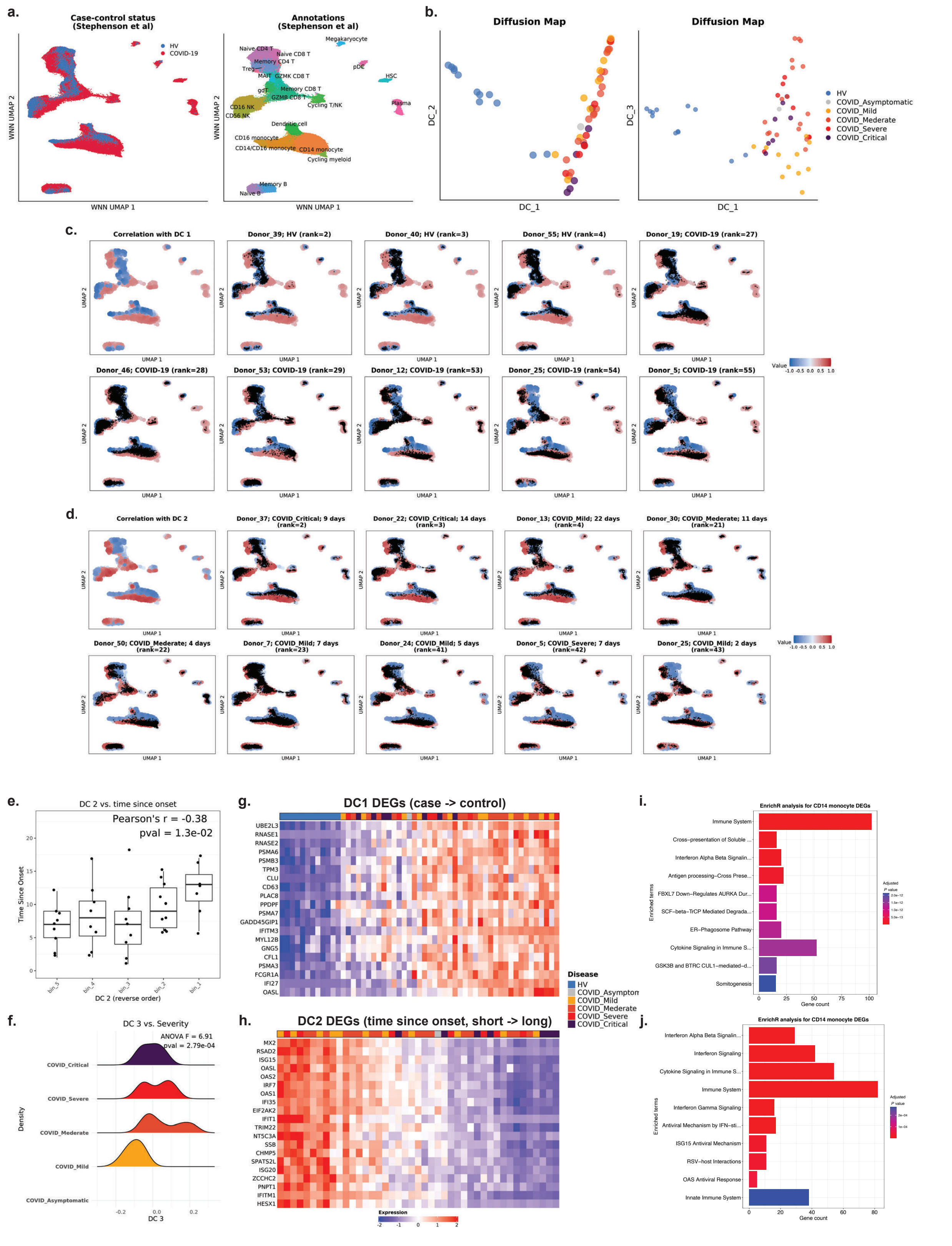


**Supplementary Figure 4. scSLIDE applied to an independent COVID-19 dataset (Stephenson et al Nat Med 2021).** (a). UMAP of cell-level weighted nearest neighbor (WNN)

embeddings from the Stephenson et al. COVID-19 dataset after scSLIDE processing. Cells colored by case–control status (left panel; HV, healthy volunteers; COVID-19, infected donors). Cells colored by major immune cell annotations (right panel). (b). Diffusion map embedding of the sample-level relative density matrix identifies principal diffusion components (DCs), with DC1 separating cases from controls and DC2 capturing heterogeneity among infected donors. (c,d). Visualizing the density of cells from each individual sample. Each colored point represents a landmark, and is colored by Pearson correlation (red, positive; blue, negative) with DC1 (c) and DC2 (d). Cells from 9 representative donors (black points) are overlaid onto the landmark map, illustrating how donor cells are distributed from samples across the DC1 and DC2 gradient, respectively. (e). DC2 correlates strongly with time since symptom onset (TSO). (f). DC3 shows a weaker severity gradient in this smaller cohort. (g,h). Heatmaps of representative differentially expressed (DE) genes whose donor-level pseudobulk expression correlates with DC1 and DC2 in CD14 monocytes, ordered by diffusion component score. (i.j). EnrichR pathway analyses for DC-associated differentially expressed genes (DEGs). The gene ontology databases being used is REACTOME (10.1093/nar/gkad1025).

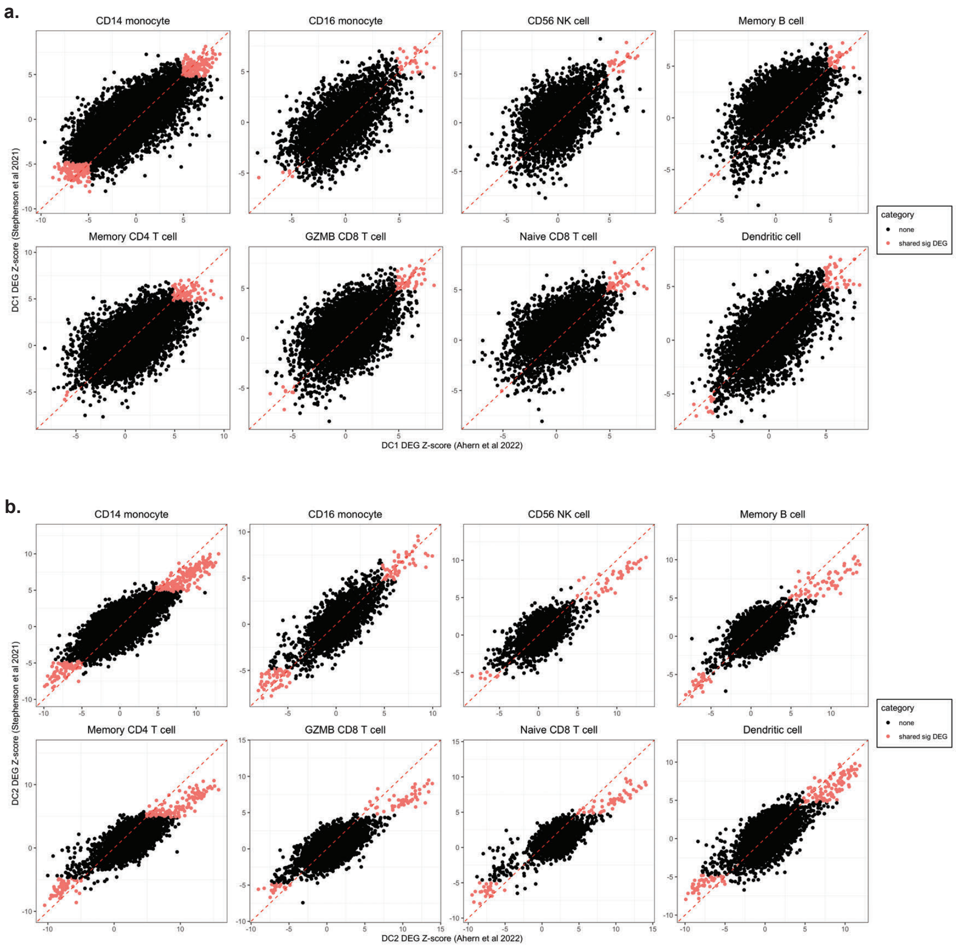


**Supplementary Figure 5.** (a–b) Correlations of DC1-associated (a) and DC2-associated (b) marker genes between the COMBAT and Stephenson datasets. Each point represents a gene's DE test z-scores from the trajectory-based NB-GLM model as independently calculated in the COMBAT dataset (x-axis) and the Stephenson dataset (y-axis). NB-GLM model was run separately for each cell type. Strong positive correlations indicate that infection-related (DC1) and interferon-related (DC2) transcriptional programs replicate across cohorts.

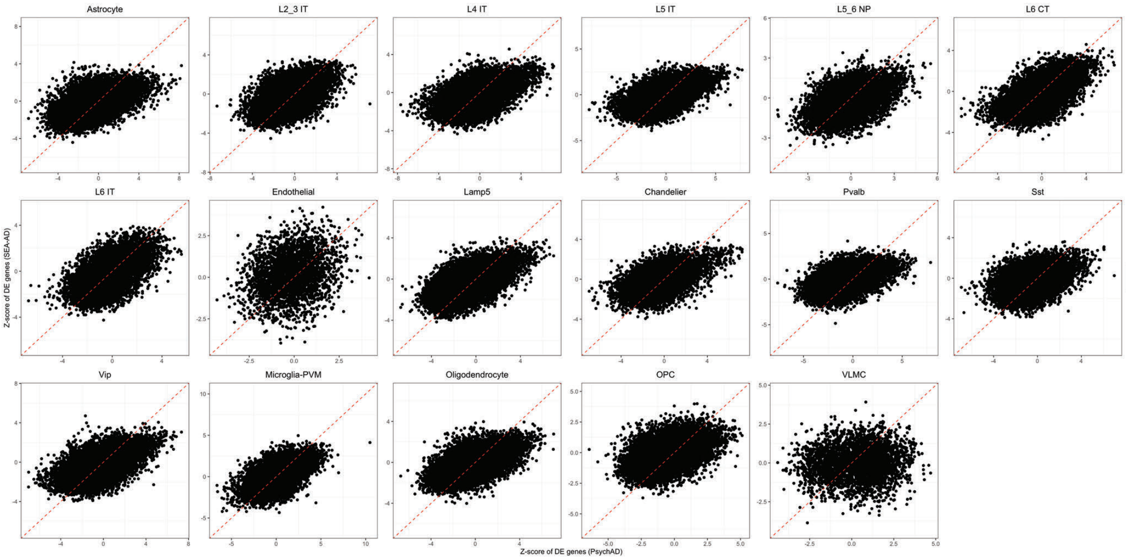


**Supplementary Figure 6.** Standard case–control pseudobulk DE analysis shows poor reproducibility between the Psych-AD and SEA-AD datasets. For each of 17 cell types, donor-level pseudobulk counts were generated and analyzed using the NB-GLM model (Supplementary Methods), modeling diagnosis as a binary case/control variable. The significance of the case/control coefficient reflects differential expression under this model. Although the analysis was performed independently for each dataset, only a small fraction of DEGs replicated across cohorts, highlighting limited cross-study consistency. We observed substantially higher reproducibility when using a continuous trajectory score instead of a binary label (Supplementary Figure 8).

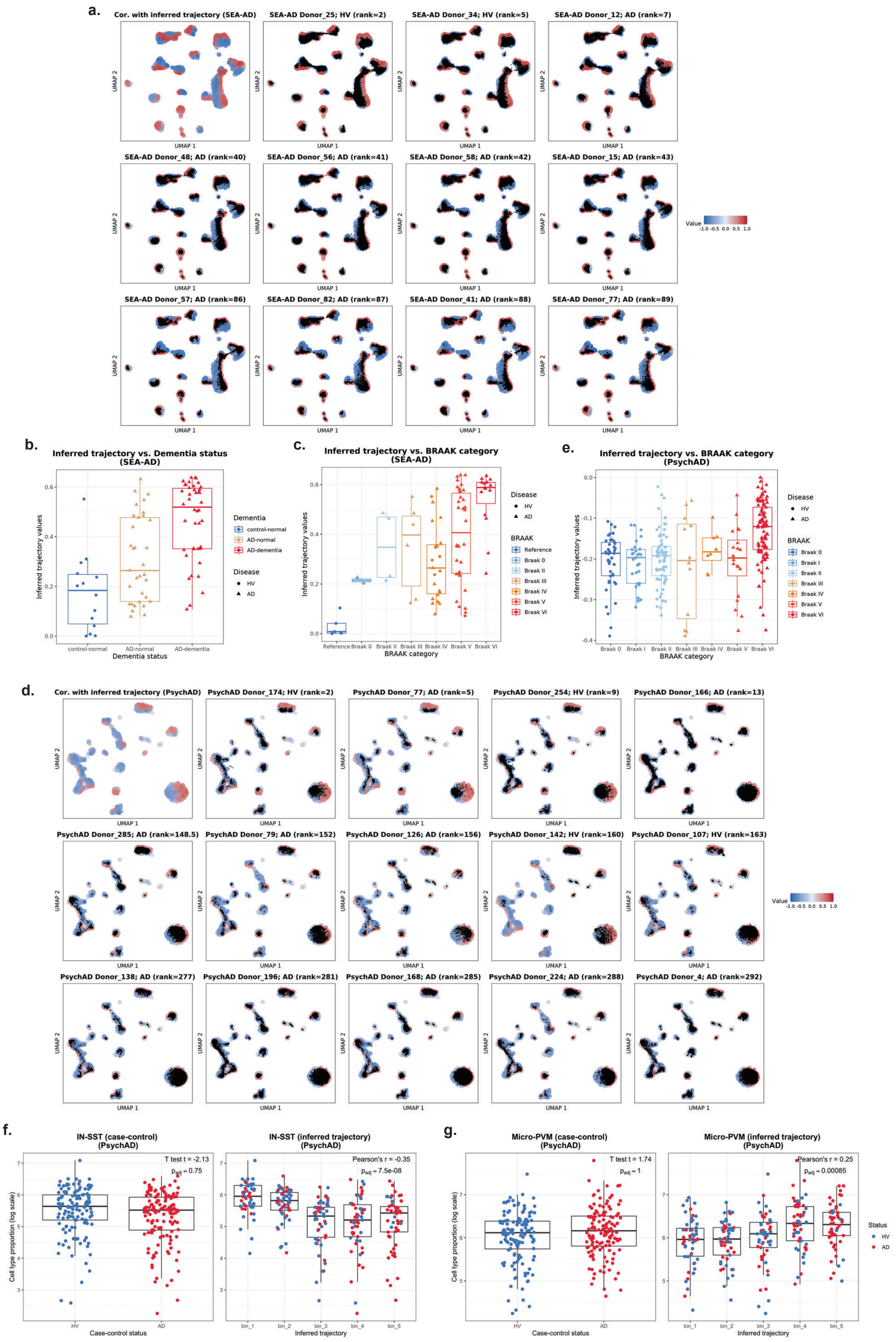

a.
Cor. with inferred trajectory (SEA-AD)
SEA-AD Donor_25; HV (rank=2)
SEA-AD Donor_34; HV (rank=5)
SEA-AD Donor_12; AD (rank=7)
SEA-AD Donor_48; AD (rank=40)
SEA-AD Donor_56; AD (rank=41)
SEA-AD Donor_58; AD (rank=42)
SEA-AD Donor_15; AD (rank=43)
SEA-AD Donor_57; AD (rank=86)
SEA-AD Donor_82; AD (rank=87)
SEA-AD Donor_41; AD (rank=88)
SEA-AD Donor_77; AD (rank=89)
UMAP 1
UMAP 2
Value
b.
Inferred trajectory vs. Dementia status (SEA-AD)
Inferred trajectory values
Dementia status
control-normal
AD-normal
AD-dementia
Dementia
Disease
HV
AD
c.
Inferred trajectory vs. BRAAK category (SEA-AD)
BRAAK category
Reference
Braak 0
Braak II
Braak III
Braak IV
Braak V
Braak VI
BRAAK
e.
Inferred trajectory vs. BRAAK category (PsychAD)
Braak I
d.
Cor. with inferred trajectory (PsychAD)
PsychAD Donor_174; HV (rank=2)
PsychAD Donor_77; AD (rank=5)
PsychAD Donor_254; HV (rank=9)
PsychAD Donor_166; AD (rank=13)
PsychAD Donor_285; AD (rank=148.5)
PsychAD Donor_79; AD (rank=152)
PsychAD Donor_126; AD (rank=156)
PsychAD Donor_142; HV (rank=160)
PsychAD Donor_107; HV (rank=163)
PsychAD Donor_138; AD (rank=277)
PsychAD Donor_196; AD (rank=281)
PsychAD Donor_168; AD (rank=285)
PsychAD Donor_224; AD (rank=288)
PsychAD Donor_4; AD (rank=292)
f.
IN-SST (case-control) (PsychAD)
T test t = -2.13
p_adj = 0.75
IN-SST (inferred trajectory) (PsychAD)
Pearson's r = -0.35
p_adj = 7.5e-08
Cell type proportion (log scale)
Case-control status
Inferred trajectory
bin_1
bin_2
bin_3
bin_4
bin_5
g.
Micro-PVM (case-control) (PsychAD)
T test t = 1.74
p_adj = 1
Micro-PVM (inferred trajectory) (PsychAD)
Pearson's r = 0.25
p_adj = 0.00085
Status

**Supplementary Figure 7.** (a). Visualizing the density of cells from each individual sample. Each colored point represents a landmark, and is colored by Pearson correlation (red, positive; blue, negative) with the inferred AD trajectory, with 11 representative donors from the SEA-AD dataset overlaid, illustrating AD variability along the inferred pseudo-trajectory. (b-c). SEA-AD donor positions along the trajectory stratified by dementia status and BRAAK category confirm association with different AD pathological measurements. (d). Same as (a) but for the Pysch-AD dataset, with 14 representative donors overlaid. (e). Psych-AD donor positions along the trajectory stratified by BRAAK category. (f,g). Cell-type abundance shifts along the inferred trajectory in Psych-AD data compared with binary case–control analysis. *SST*+ interneurons (h) decrease, and microglia (i) increase progressively along the trajectory, consistent what was observed in SEA-AD data shown in Figure 3h,i.

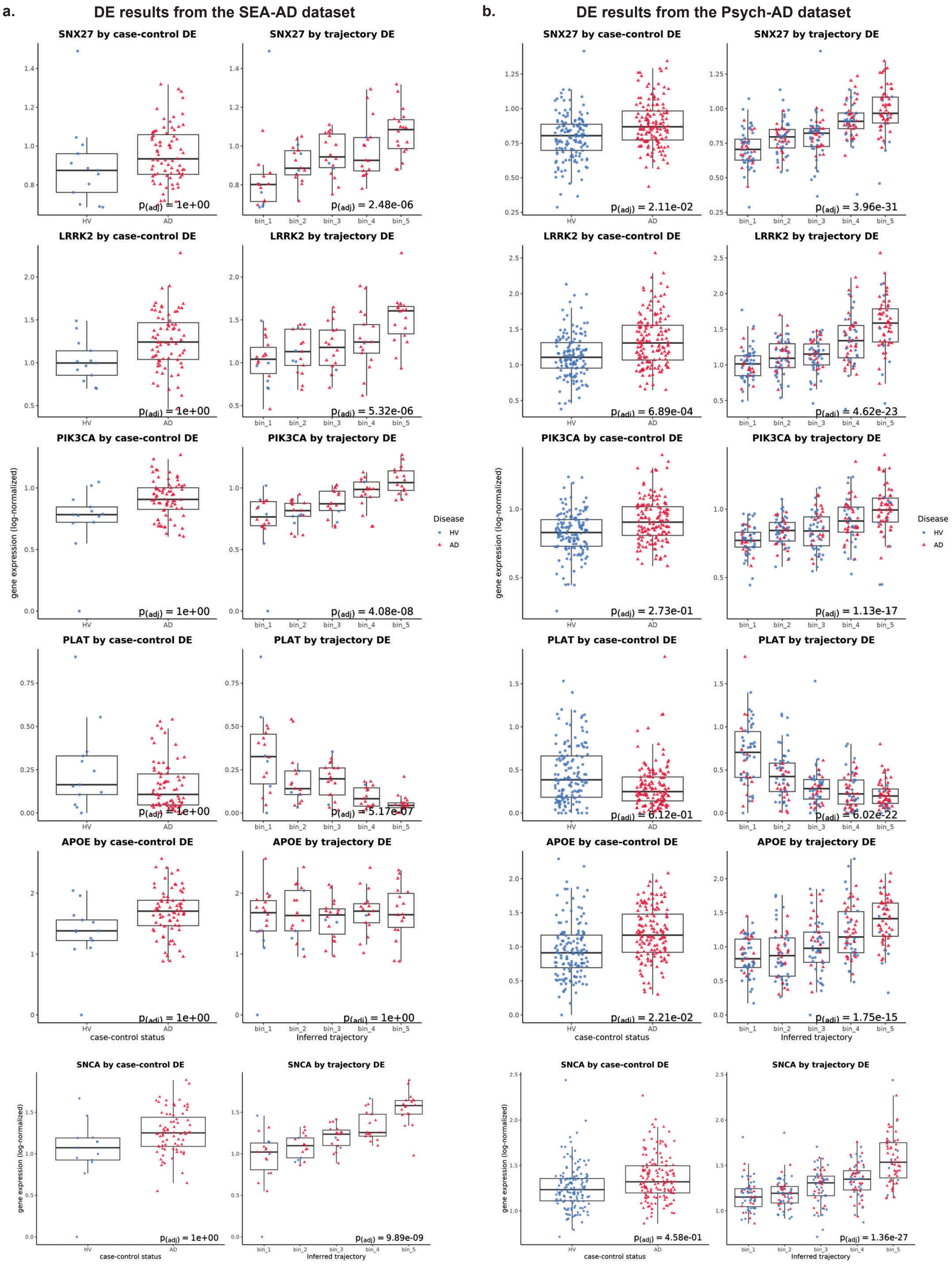


**Supplementary Figure 8.** (a–b) Expression of canonical and genetically/biologically implicated AD genes (e.g., *SNCA, LRRK2, SNX27, PIK3CA, PLAT, APOE*) in the SEA-AD and Psych-AD datasets. (a) Shows expression patterns for SEA-AD, and (b) shows pattern for Psych-AD. With each panel left boxplot shows the pseudobulked expression (per-donor; within microglia), split by binary case/control status. In most cases, the genes show weak statistical evidence of differential expression that does not survive multiple testing correction. Right boxplot shows the expression patterns with samples grouped into five equally sized bins based on their inferred position in the disease trajectory. Grouping by disease progression substantially increases statistical power to identify changes in both datasets. We observe statistically significant changes (after multiple testing correction) that replicate in both datasets in all cases, with the exception of *APOE*.

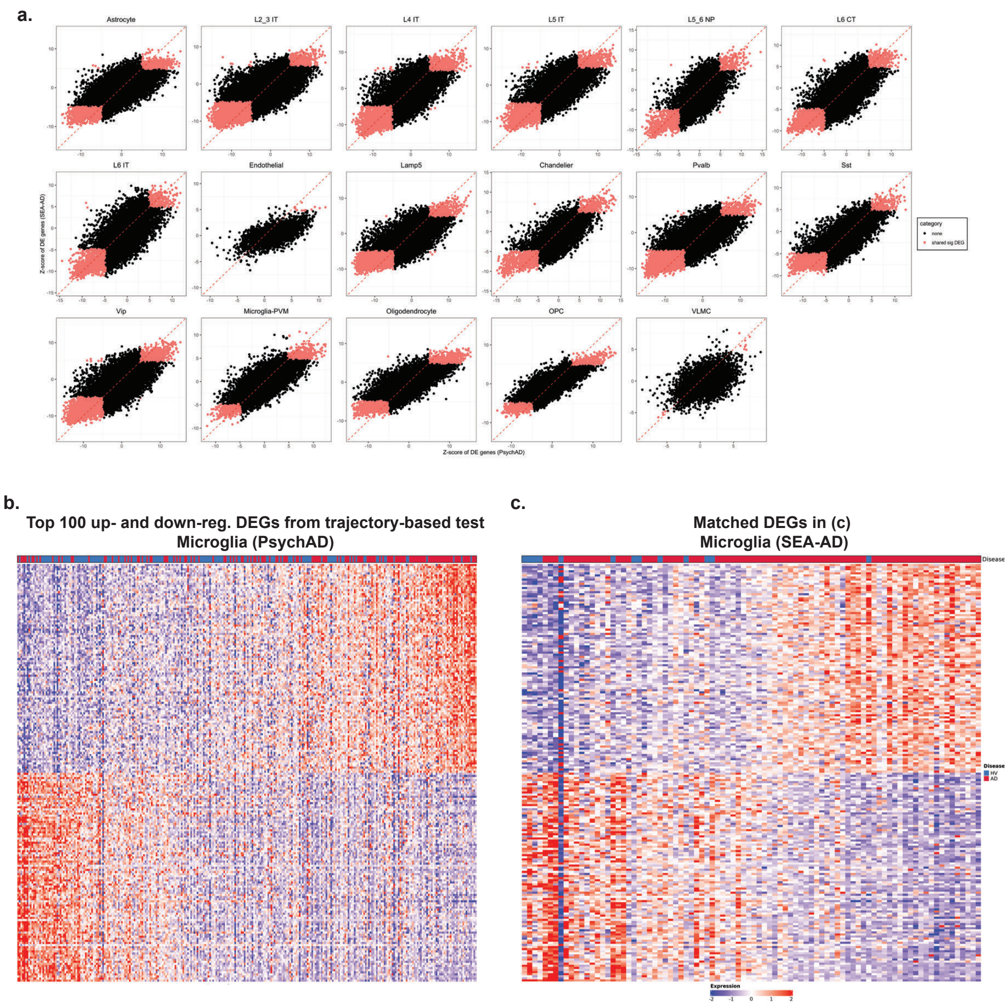


**Supplementary Figure 9.** (a). Scatterplots comparing per-gene trajectory-based DE test z-scores between SEA-AD and Psych-AD datasets, showing high concordance for AD DE gens across multiple cell types. (b). Heatmaps of top 100 trajectory-associated DEGs in microglia identified in Psych-AD (b) and matched DEGs in SEA-AD (c) showing highly reproducible transcriptional signatures. The same genes (which were first identified by NB-GLM analysis of the Psych-AD dataset, and then shown to independently replicate in the SEA-AD dataset) are shown in both heatmaps, and in the same order.

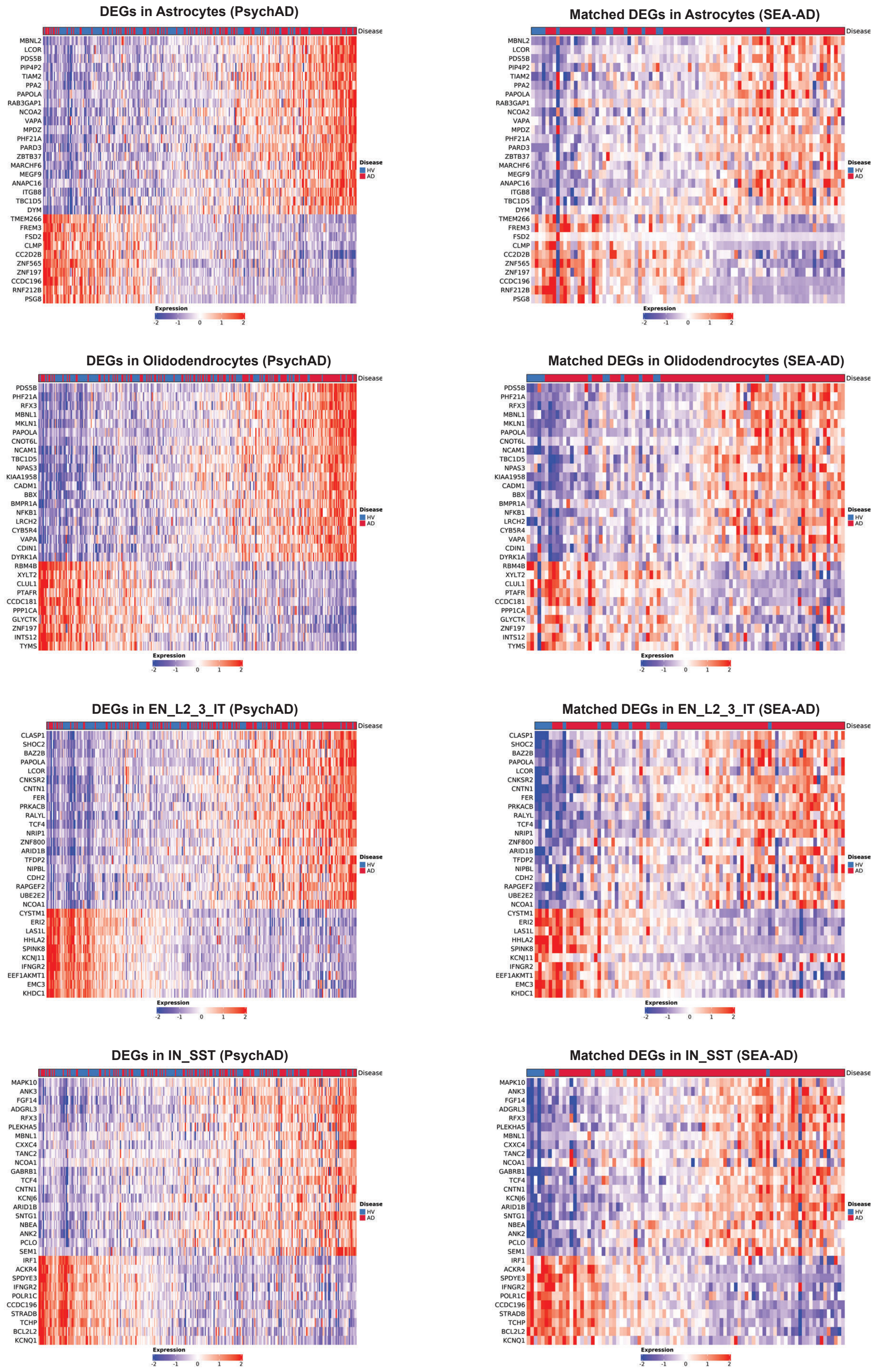


**Supplementary Figure 10.** Paired comparisons of trajectory-associated DEGs across Psych-AD and SEA-AD for multiple cell types, including astrocytes, oligodendrocytes, excitatory

EN_L2_3_IT neurons, and inhibitory IN_SST neurons (*SST*+ interneurons), demonstrate cross-cell type concordance. The same genes (which were first identified by NB-GLM analysis of the Psych-AD dataset, and then shown to independently replicate in the SEA-AD dataset) are shown in both heatmaps, and in the same order.

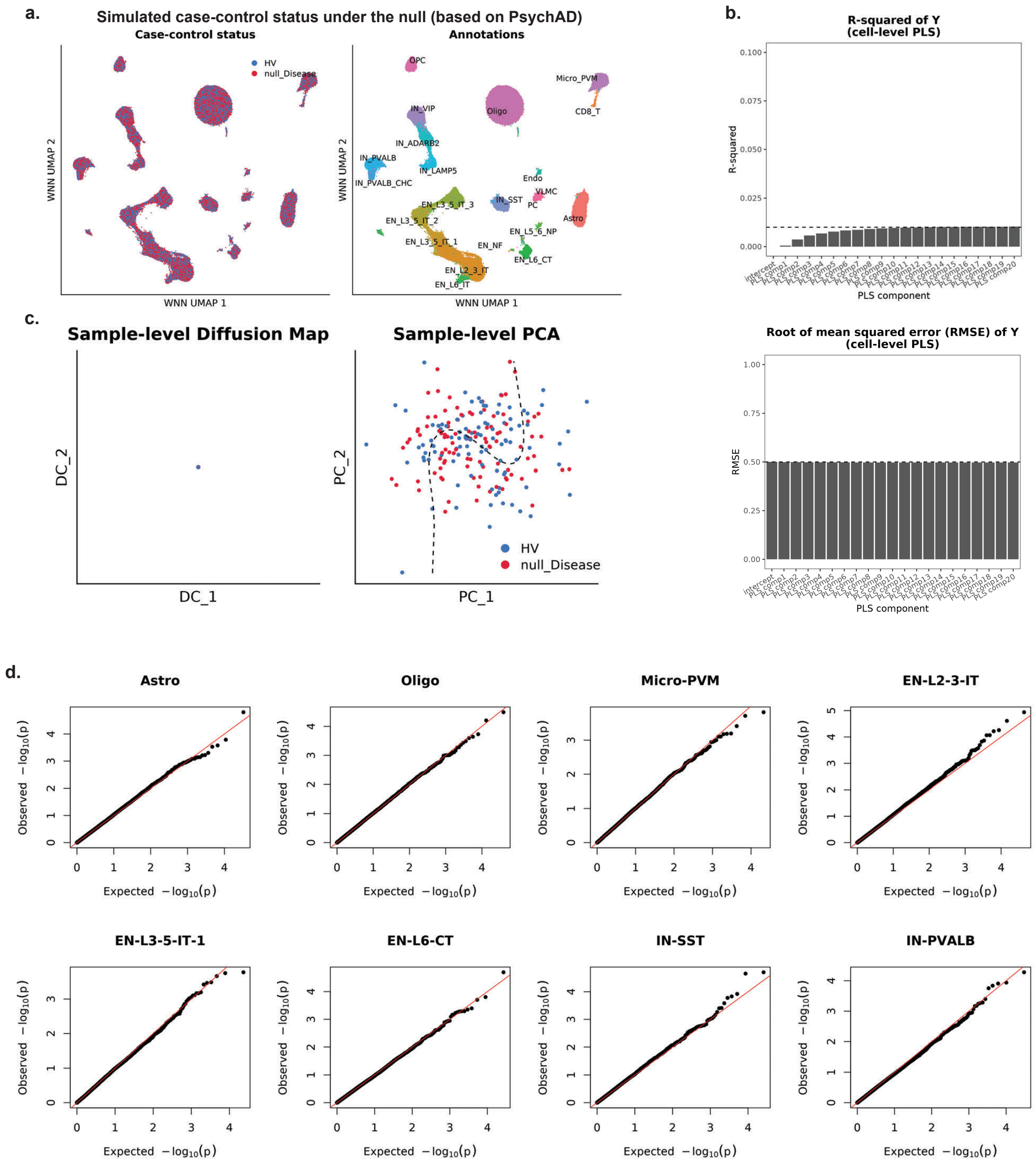


**Supplementary Figure 11. Null simulations demonstrate specificity of scSLIDE.** (a). Cell-level WNN UMAP for simulated data under the null scenario. (b). The $r^2$ and root of mean squared error (RMSE) of cell-level partial least squares analysis showing no separation between simulated case-control labels under the null. (c). Sample-level embeddings (diffusion map and principal component analysis) also show no separation between simulated case-control labels under the null. (d). No DEGs are detected along the fitted principal curve, indicating an absence of spurious structure.

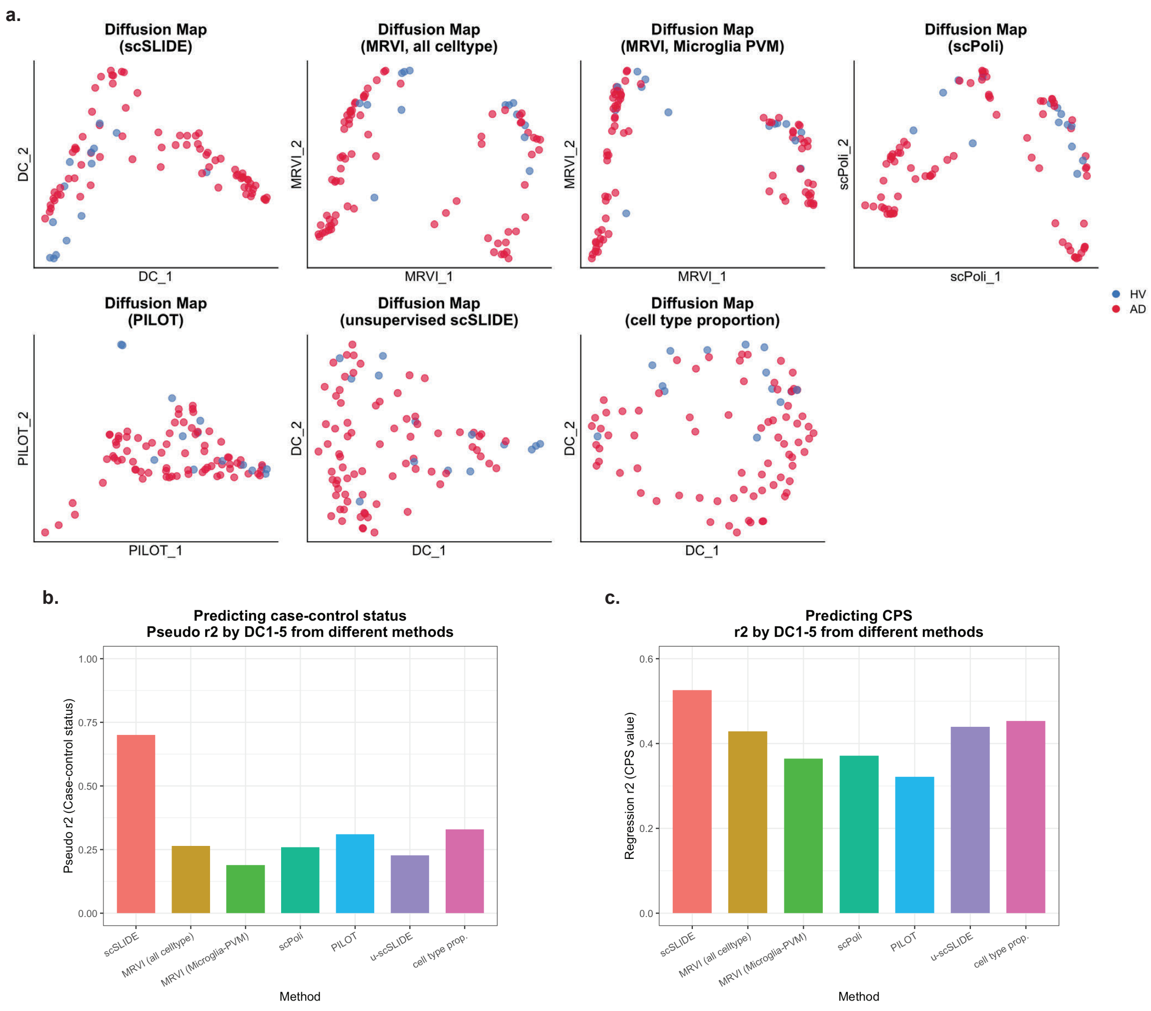


**Supplementary Figure 12. Benchmarking on the SEA-AD data using alternative sample-level methods.** (a) Diffusion map embeddings derived from scSLIDE, MRVI (based on all cell types), MRVI (based on microglia PVM), scPoli, PILOT, an unsupervised scSLIDE variant (u-scSLIDE, see Supplementary Methods), and a cell-type-proportion workflow. Each point is colored based on each individual's case-control status. (b). To assess how well each method recapitulates case-control status, we fitted logistic regression models using the top five diffusion components (DCs) as predictors. The bar plot shows McFadden's pseudo-$r^2$ for each model, comparing predictive performance across methods. (c). Similar to (b), but here we fitted linear regression models using the top five DCs to predict the CPS values instead. The bar plot shows the regression $r^2$ for each model.

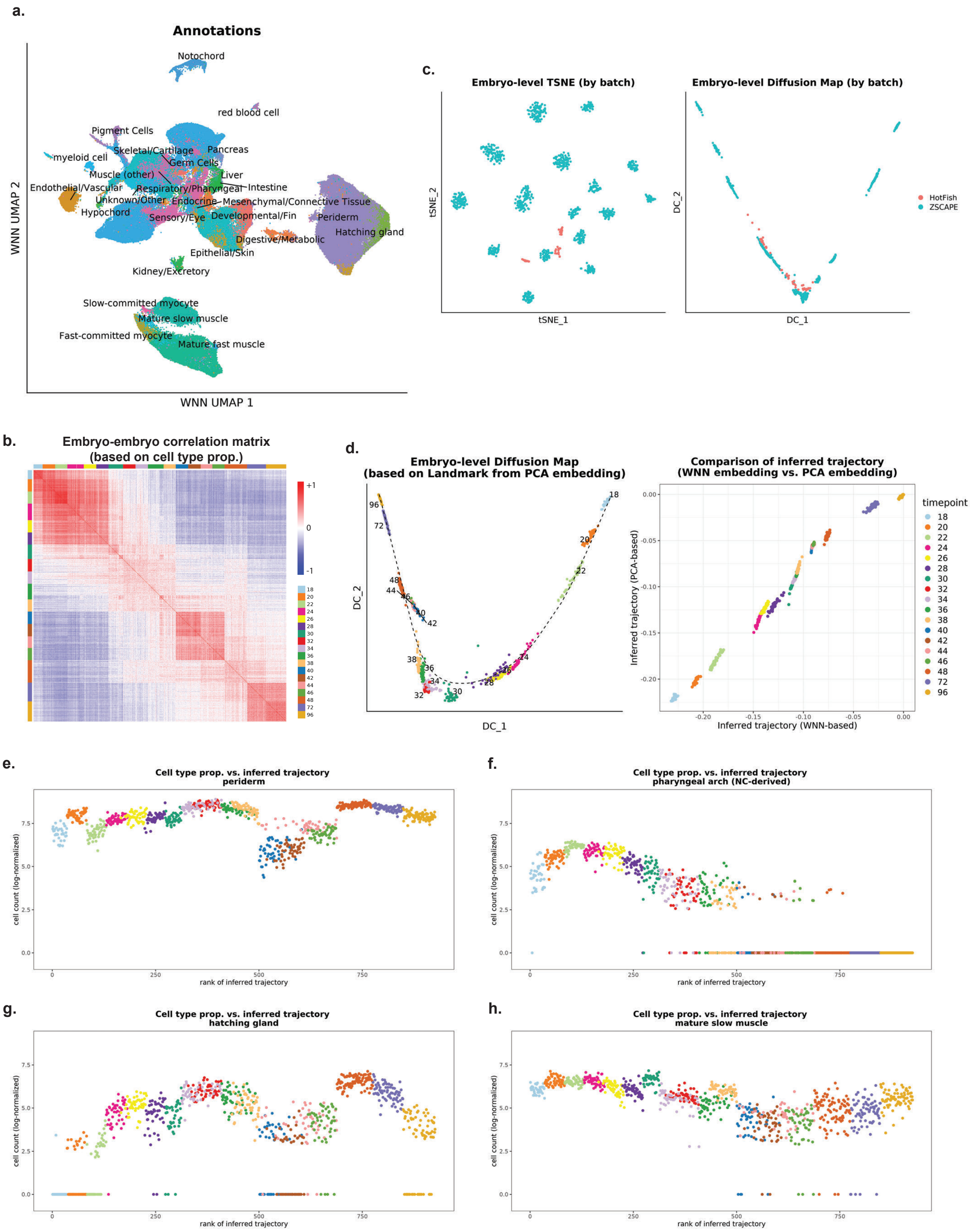


**Supplementary Figure 13.** (a). WNN cell-level embedding of the zebrafish embryogenesis dataset, with cells colored by cell type annotations. (b). Embryo-embryo correlation matrix calculated based on the normalized embryo-level cell type proportion matrix. (c). Embryo-level t-SNE embedding (left) and diffusion map embedding (right) reveal outlier embryos, which were subsequently found to be grown at different temperatures. (d). Fully unsupervised scSLIDE (without PLS) reconstructs a consistent developmental trajectory that is consistent with standard scSLIDE analysis. Left panel: the diffusion map of embryo-level density matrix from unsupervised scSLIDE; right panel: the comparison of inferred trajectories between standard scSLIDE and unsupervised scSLIDE. (e-h). Additional examples (as shown in Figure 5f) of cell types whose dynamic abundance varies both across and within developmental timepoints. Embryos are ordered by their trajectory-inferred progression, revealing gradual transitions in the abundance of multiple cell types (periderm, pharyngeal arch, hatching gland, mature slow muscle).
